# Evolutionary metabolomics of specialized metabolism diversification in the genus *Nicotiana* highlights allopolyploidy-mediated innovations in *N*-acylnornicotine metabolism

**DOI:** 10.1101/2022.09.12.507566

**Authors:** David Elser, David Pflieger, Claire Villette, Baptiste Moegle, Laurence Miesch, Emmanuel Gaquerel

## Abstract

Specialized metabolite (SM) diversification is a core process to plants’ adaptation to diverse ecological niches. Here we implemented a computational mass spectrometry (MS)-based metabolomics approach to explore SM diversification in tissues of 20 species covering *Nicotiana* phylogenetics sections. To drastically increase metabolite annotation, we created a large *in silico* fragmentation database, comprising more than 1 million structures, and scripts for connecting class prediction to consensus substructures. Altogether, the approach provides an unprecedented cartography of SM diversity and section-specific innovations in this genus. As a case-study, and in combination with NMR and MS imaging, we explored the distribution of *N-*acyl nornicotines, alkaloids predicted to be specific to *Repandae* allopolyploids, and revealed their prevalence in the genus, albeit at much lower magnitude, as well as a greater structural diversity than previously thought. Altogether, the novel data integration approaches provided here should act as a resource for future research in plant SM evolution.

**Teaser:** Computational metabolomics delineates main trends in the diversification of specialized metabolism in the genus *Nicotiana*

## Introduction

Plant metabolic profiles represent complex traits that reflect both evolutionary and temporally dynamic adaptations to specific ecological niches. Compared with their counterparts integrated into broadly conserved central C metabolism pathways, specialized metabolites (SMs) contribute to the largest fraction of inter-specific variations in plant metabolic profiles. This plant chemodiversity is predicted to account for somewhere on the order of one hundred thousand to one million chemically unique structures, with an estimated range of five thousand to fifteen thousand structures *per* plant species (*1*). Many of these SMs act as chemical shields or as attractants in plant biotic interactions. In this respect, a relatively recent paradigm shift as part of ecological hypotheses such as the synergy (*2*) and interaction diversity hypotheses (*3*), has been to consider SM structural diversity, and not solely the summation of individual metabolites, as a critical determinant of plants’ ecological interactions. The latter perspective also revives the interest in the exploration of SM diversity with modern analytical approaches and the use of adequate statistical descriptors (*4*).

In analogy to phylogenomics approaches that have flourished as a result of both the increasing release of annotated genomes and of established comparative bioinformatics pipelines to analyze these data, recent years have indeed seen a resurgence of plant family-/genus-specific comparative metabolomics analyses to guide functional biochemical studies. For instance, comparative metabolomics within the Rhamnaceae revealed that only the Ziziphoid clade of this family possesses a functional triterpenoid biosynthetic pathway, whereas the Rhamnoid clade predominantly developed diversity in flavonoid glycosides (*5*). In a previous study, we implemented a metabolomics-centered fragmentation rule-based pipeline to annotate the diversity of 17-hydroxygeranyl linalool (17-HGL) diterpene glycosides within the Solanaceae family and revealed intense chemotypic structural variations combined with a patchy distribution of this compound class as it appeared restricted to the *Capsicum, Lycium* and *Nicotiana* genera (*6*). The latter “phylometabolomics” information facilitated gene candidate selection for functional biochemical studies in the 17-HGL diterpene glycoside pathway (*6*). Similarly, comparative metabolomics across multiple Solanaceae species was instrumental in guiding co-expression studies for gene discovery for the steroidal glycoalkaloid pathway emblematic of the *Solanum* genus (*7*).

Besides inter-specific variations in SMs, another important dimension, unfortunately rarely integrated into taxonomic-scale metabolomics studies, is the tissue/organ specialization of most SM pathways. Exploring these tissue/organ-level variations and their statistical correlation with gene expression data can be extremely powerful in the process of SM biosynthetic gene discovery (*8*). Analyzing tissue-specific metabolomes is also critical to test ecological theories of plant investments into metabolic defenses such as the optimal defense theory which predicts greater metabolic defense accumulation in developmental stages/tissues with higher organismic-level fitness contribution and/or greater predation rates (*9*). Trichomes, in particular glandular ones covering most aerial plant tissues, are notorious for their capacity to synthesize high amounts of very specific SMs (*10*). Trichome SMs can be either stored within trichome cells and glands or actively secreted, such as for Solanaceae-specific poly-acylated sugars, also referred to as *O*-acyl sugars and whose biochemistry has been thoroughly investigated in recent years (*11*). Calyces formed by the floral sepals and which protect maturating reproductive organs are typically rich in SMs whose biosynthesis can be dependent on trichomes present on these tissues (*12–14*). SM profiles of roots, while much less systematically explored than shoot-based ones, can be as structurally diverse as trichome-specific ones (*15*) and have recently become of major focus for our understanding of SM ecological functions for plant-soil microorganisms’ interactions (*16*).

Most recent advances in computational metabolomics provide a long-awaited framework to systematically explore the above-described importance of the species *x* tissue SM variations (*4*). These novel capacities to explore plant chemical spaces are further propelled by platforms such as, the MassIVE database (https://massive.ucsd.edu/) reaching 12000 metabolomics datasets in 2022. Despite the increasing amount of data that can be generated from modern MS instruments, the average annotation rate of most MS metabolomics studies remains at the order of a few percent of deconvoluted MS/MS features (*17*). The number of computational tools to address this challenge of transforming spectral information into chemical knowledge is hence rapidly increasing and can be divided into two main approaches. One set of approaches relies on MS/MS spectral grouping, as embodied by the game-changing development of molecular networking and of the repertoire of network annotation/mining tools embedded within the GNPS ecosystem (*18, 19*). A second set of approaches relies on *in silico* fragmentation and (sub)structure prediction from mass spectra. Classification of spectra within ontologies of molecular families can notably be achieved by CANOPUS, a deep neural network method which is able to predict 2497 compound classes (*20*) and which is embedded with the elemental formula prediction and structure annotation pipeline from SIRIUS (*21*). Alternatively, the MS2LDA method allows to extract information derived from shared substructures from spectral data via a Latent Dirichlet Allocation algorithm borrowed from topic modelling (*22*). Recently developed or significantly upgraded computational tools such as CFM-ID (*23*), molDiscovery (*24*), Metfrag (*25*) or QCxMS (*26*) provide algorithmic means to predict MS/MS spectra from structures. However, systematically prioritizing and/or merging the highest confidence predictions from each of these tools remains a challenge that is rarely tackled in most MS metabolomic studies.

The genus *Nicotiana L*. combines several appealing features to study SM pathway diversification. This genus, comprising 13 well phylogenetically-resolved sections for a total of at least 80 species, is appearing in various morphological forms such as small herbs to shrubs up to small trees, which often are viscid-glandular and rarely glabrous (*27, 28*). Among the most studied species in this genus are *Nicotiana tabacum* and *N. rustica*, which are traditionally grown for tobacco products; *N. glauca*, which has been a focus of biofuel research studies (*29*) and *N. benthamiana*, a very popular model organism in molecular biology (*30*). The intense research on *Nicotiana* species is further reflected into the very large set of reference transcriptome and genome resources publicly available for species of this genus (*31*). As recently reviewed (*32*), the phytochemistry of several species of this genus, in particular that of the coyote tobacco *Nicotiana attenuata*, a flagship model organism for the chemical ecology of plant-insect interactions (*33*), has been extensively studied with notable focus on alkaloids, mono/sesquiterpene volatiles, *O*-acyl sugars or 17-HGL diterpene glycosides. Finally, half of the species of the *Nicotiana* genus are allopolyploids of different ages and for some of them, the closest extant diploid progenitors have been mapped, thereby providing a phylogenetics framework to study allopolyploidy-mediated contributions to phenotypic trait evolution (*34*). Among the *Nicotiana* SM innovations thought to have been shaped by recent allopolyploidization events are *N-*acyl-nornicotines (NANNs), derived from the *N*-acylation of nornicotine with long chain fatty acyl chains and which have been described as specific to allopolyploid species of the section *Repandae* (*35*). This *Nicotiana* section is about 4.5 million years old and has *N. sylvestris* as its closest diploid maternal and *N. obtusifolia* as closest paternal progenitor. The 17 NANN structures originally described in the *Repandae* species *N. repanda, N. nesophila, N. stocktonii* (*36, 37*), but not in *N. nudicaulis*, likely act as gain-of-function anti-herbivory defenses. Indeed, compared with the *Nicotiana* widespread nicotine and nornicotine non-acylated alkaloids, NANNs are highly effective against and evade the resistance acquired for nicotine/nornicotine by the tobacco hornworm *Manduca sexta*, a native herbivore (*38*). However, the evolution of this defensive trait is largely underexplored and detailed investigations on the NANN structural diversity within the genus *Nicotiana* are missing.

Here, we implemented a comprehensive computational MS metabolomics workflow to explore SM chemodiversity in various tissues of 20 species representative of the main phylogenetic sections within the *Nicotiana* genus. By employing a multi-inference deep annotation approach that ultimately connects information theory statistics, chemical class mapping and substructure inferences, we provide an unprecedented cartography of SM tissue-level distribution in this genus. The results of this study provide access to novel SM annotations and tissue *x* species distribution data to guide future biochemical studies and notably shed light on the unsuspected structural diversity and evolutionary trajectory of the NANNs defensive pathway within the *Nicotiana* genus.

## Results

### Tissue-level metabolomics data capture phylogenetically-relevant Nicotiana SM diversity

In order to comprehensively explore tissue-level SM diversification in the *Nicotiana* genus, we profiled the metabolome of leaves elicited or not with methyljasmonate, concentrated leaf surface exudates, complete root and of calyces (**Fig. 1B**) of 20 species covering all of the main sections of this genus as well as diploid and allotetraploid states (**Fig.1A** and **Table S1**). Besides phylogenetic position, species selection further took into consideration the availability of transcriptomics/genomics data as a platform for future functional studies (*31*). Sampled tissues were selected based on previous studies of our group (*8*) indicating the high degree of tissue-level specialization in SM distribution and conversely the importance of concatenating multi-tissue profiles to increase SM coverage. Additionally, we aimed via this pluri-tissue approach to explore tissue-level shifts in SM class prevalence across the focal species as a mechanism of organismic-level chemodiversification. Noteworthy, amounts of leaf exudate material collected greatly differed among the focal species, with *Nicotiana setchellii* (2.7 mg of exudate *per* g leaf fresh weight) and *Nicotiana glutinosa* (2.5 mg/g) producing the largest amounts of dried exudates collected from leaf washes (**Fig. S1** and **Table S1**). All methanolic extracts were analyzed using a previously established UPLC-ESI^+^QTOFMS method with optimized settings for massive MS/MS data collection (*6*). 17901 metabolite-derived MS/MS spectra (hereafter referred to as features) were, after a data redundancy and contaminant check using a custom script, deconvoluted and considered for Feature-based Molecular Networking (FBMN) processing with settings that were optimized to handle the species *x* tissue-exacerbated metabolic diversity in the dataset. The resulting species *x* tissue MS/MS feature compendium served as input for the data exploration workflow presented in **Fig. 1D** (**Fig. S2**).

**Figure 1.**
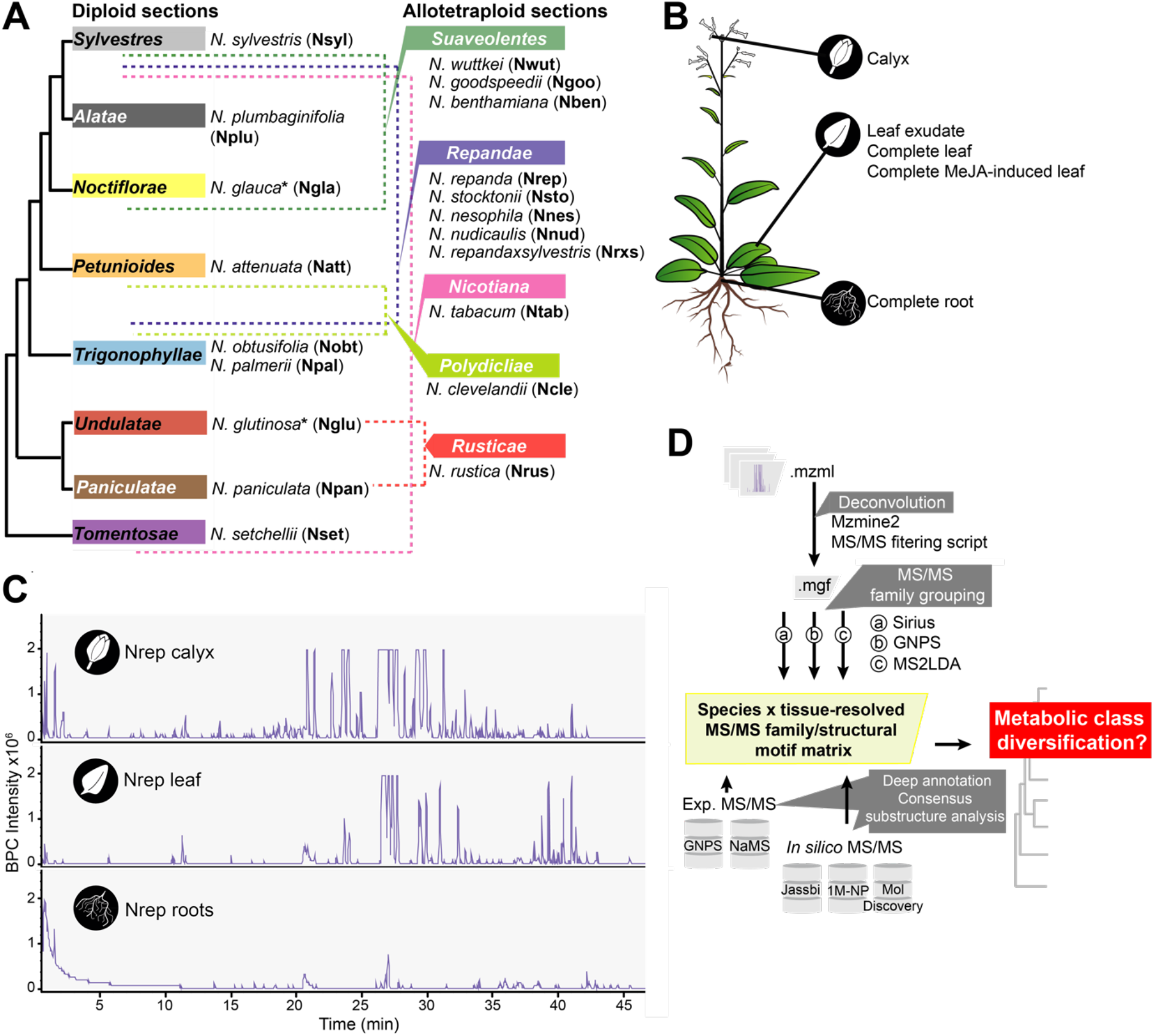
Experimental and data processing set-ups to explore species *x* tissue specialized metabolism diversification in the *Nicotiana* genus. **(A)** Schematic *Nicotiana* phylogenetic tree highlighting main genus sections and representative species selected for metabolomics analysis. Four allotretraploid sections, dashed lines indicate sections containing closest extant diploid progenitors. Accessions and origins of the selected species are referred to in **Table S1**. *N. glutinosa*\* and *N. glauca*\* are considered as homoploid hybrids as summarized in (*34*). **(B)** Tissue sampled from 6-to-8 weeks old plants of the selected 20 *Nicotiana* species. Fully elongated leaves were considered for leaf-based samplings. Leaf exudates were prepared by acetonitrile-based leaf surface rinsing; methyljasmonate (MeJA)-treated leaves were harvested 72h post-treatment. **(C)** Representative Base Peak Chromatograms (BPC) from the UPLC-ESI^+^-QTOF MS analysis of methanolic extracts of *N. repanda* roots, untreated leaves and calyces. **(D)** Data processing pipeline to construct a species *x* tissue MS/MS spectral matrix and for its deep structural annotation prior to metabolic class diversification analysis. Architecture of the data processing pipeline and organization of the different output matrices as supplementary data-sets are presented in **Fig. S2**.

To contrast patterns of feature diversity across species, we calculated, for each of the tissue types, α-diversity scores based on the Shannon Entropy (*H*) from Information Theory (*8*) (**Fig. 2A**). A unifying trend in these tissue-level analyses was that species’ profiles, differed extremely in their α-diversity indices, up to 3-fold counter-species variations being detected depending on the tissue type. Root samples were, from all examined plant samples, those with consistently lower α-diversity scores (average *H* = 7.6), likely indicative of the prevalence of only a few SM classes in these samples for the analytical conditions considered in this study (**Fig. 1C**). As expected, highest α-diversity scores were on average detected for MeJA-elicited leaves (average *H* = 9.4) (**Fig. S3**), followed by uninduced leaves (average *H* = 9.3) and calyces (average *H* = 9.0). The effect of the MeJA elicitation on feature diversity was consistently more apparent at the level of detected features and very variable among the focal species (**Fig. S3**). Interestingly, we noted that these inter-species variations in MeJA inducibility (indicative of the amplitude of a “metabolome plasticity” to this treatment) were strongly negatively correlated (Pearson Correlation Coefficient = -0.76, *P*-value = 1.04 × 10^−4^) with α-diversity scores of uninduced leaves (“constitutive diversity”) (**Fig. S3**). Additionally, while we initially assumed that the metabolic profiles of the exudates collected from uninduced leaves would be restricted to a few prevalent SMs (thereby resulting into in low α-diversity scores for this sample type), the relatively high α-diversity scores detected in most species were consistent with a far greater chemical diversity in those extracts. In clear constrast, *Repandae* species, with the exception of *N. nudicaulis* and the hybrid *N. sylvestris x N. repanda*, exhibited much lower α-diversity scores (*H* ranging from 3.5 to 3.7 compared with the average *H* value of 8.5 for the rest of the species) that were in line with the previously reported over-dominance of NANNs within their exudates (*37*). When feasible based on the species sampling, we also compared the α-diversity scores of allotetraploid species to those of closest diploid progenitors. Independently of the tissue type considered, we did not observe evidence of clear metabolic additivity in allotetraploid species, which would translate into higher α-diversity scores as compared to those of closest diploid progenitors (**Fig. 2A**).

**Figure 2.**
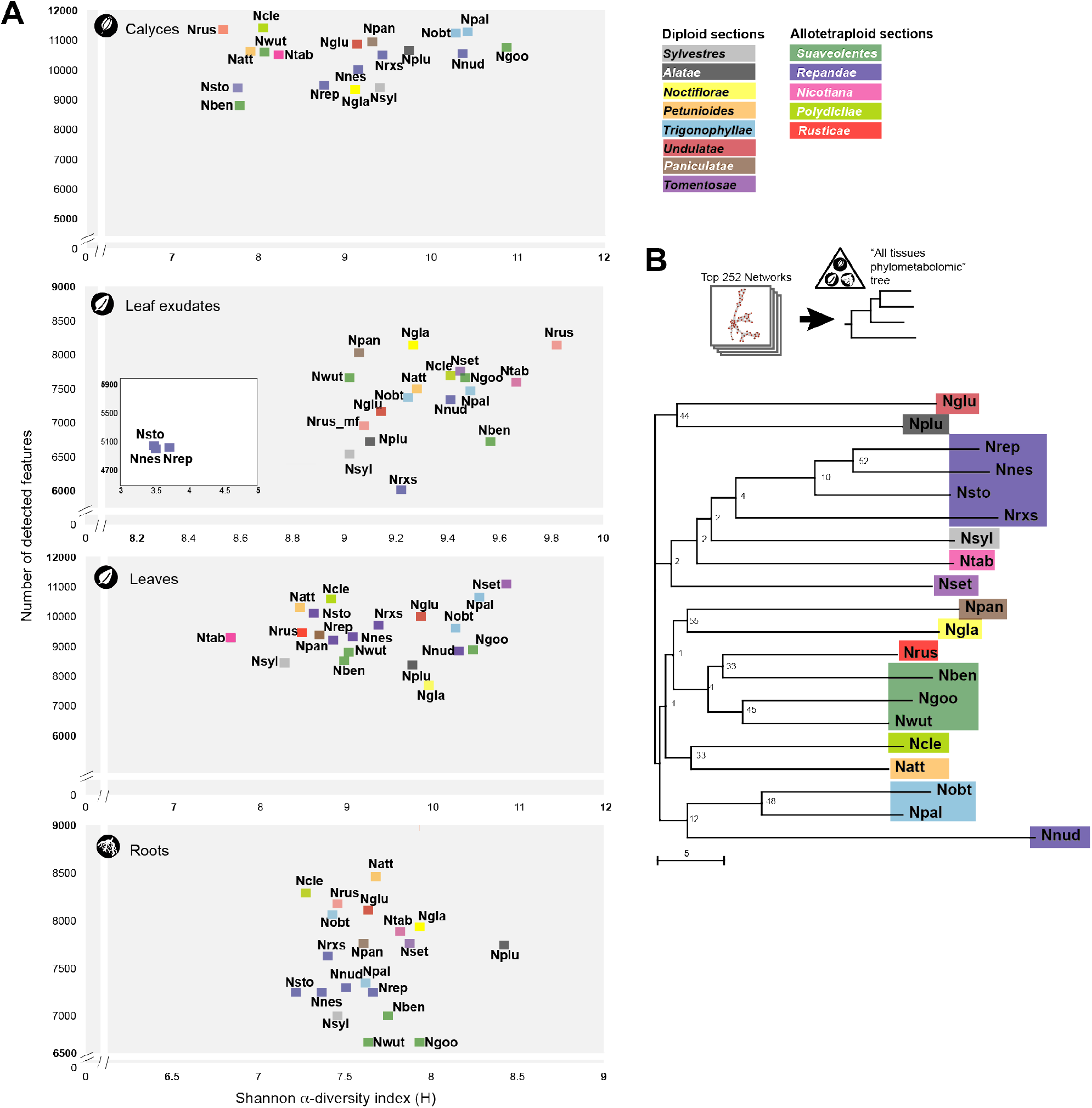
Species metabolome α-diversity and “phylometabolomics” relatedness. **(A)** Biplots depict the number of detected features and the Information Theory Shannon α-diversity as an index of feature richness *per* tissue. *Nicotiana* phylogenetic sections are color-coded. **(B)** “Phylometabolomics tree computed from the molecular networking information. To analyze the relatedness of species’ metabolomes, we first computed inter-species Euclidean distances based the molecular networking information and used the resulting distance matrices for constructing a “phylometabolomics” tree based on the Neighbor-Joining algorithm (bootstrap values derived from 999 iterations) (**Table S2**). Trees were also constructed from the tissue-level data (**Fig. S4**).

To analyze the relatedness of species’ metabolomes, we further computed inter-species metabolic distances based the molecular networking information and used the resulting distance matrices for constructing “phylometabolomics” trees. Several studies had previously attempted to construct such “phylometabolomics” trees but from single-tissue metabolome data. Here, we constructed trees both from the tissue-level (**Fig. S4**) and combined tissue data (**Fig. 2B**). The resulting “all tissues” phylometabolomics tree captured patterns of metabolome-relatedness that were frequently in accordance with the species’ tree section-level grouping and relatedness (**Fig. 2B**). Among other interesting insights, *Repandae* species’ metabolomes, with the exception of that of *N. nudicaulis*, appeared much closely-related at the “all tissues” metabolome level to that the *Sylvestres* section from which their maternal progenitor had been associated with, than to the *Trigonophyllae* section (paternal progenitor section) (**Fig. 2B, Table S2**).

### Creating a cartography of Nicotiana SM class diversification

After highlighting counter-species chemodiversity variations, we then systematically characterized onto which SM classes they mapped. In analogy to gene family inference and survey across focal species as a first step in phylogenomics, we first employed the CANOPUS tool for *ad hoc* systematic compound class and chemical ontology predictions. To combine FBMN and CANOPUS information, we implemented a frequency-based molecular <u>n</u>etwork-based <u>p</u>ropagation of CANOPUS (NP-CANOPUS) predictions, resulting into class predictions for 86.5 % of the total features within the 1586 networks retrieved by FBMN. CANOPUS “super-class” and “most specific class” intensity distributions integrating all tissue samples of given species were encapsulated as treemaps and mapped onto the species tree to provide a bird’s eye view on class expansions and shrinkages (**Fig. 3B**). For the sake of simplicity, only a few of the main tendencies are reported below; close-up views on particular “metabolic tiles” and tissue-specific treemaps are accessible in **Data S1**. Most clearly apparent was the highest proportion of “lipids and lipid-like molecules” in all species, with a significant fraction of these lipids being, in many species, contributed by the saccharolipid sub-class commonly referred to as *O*-acyl sugars in the Solanaceae. Browsing these treemaps supported the presence of high amounts of predicted diterpenes in *N. tabacum, N. sylvestris* and the cross between *N. repanda x N. sylvestris* – the latter hybrid having been initially incorporated to test progenitor chemical trait dominance. Among other trends, this analysis also pinpointed on *N. setchellii* exhibiting the most diverse and abundant set of “phenylpropanoid derivatives” from predicted 3-*O*-methylated flavonoids (connected to network #361), simple hydroxycinnamic acids (network #990), up to coumarin glycosides (network #532). Noteworthy, the performance of CANOPUS predictions was nonetheless hampered for SMs that contained substructures from independent biosynthetic origins, thereby resulting into heterogeneous CANOPUS ontologies. For instance, the large “amino acids and derivatives” tile within the *N. glauca* treemap was mostly associated with network #468, but the features embedded in this network were manually curated as *N-* hydroxycinnamoyl-spermidine conjugates which are commonly encountered in leaves of Solanaceae species as antiherbivore defenses (*39*). Also highlighting this limitation was that the high-level of NANNs which are emblematic of the *Repandae* section, was not as easily noticeable on the corresponding treemaps. Previously characterized NANNs were indeed split into several classes as “organoheterocyclic compounds”, “benzenoids” and “organic nitrogen compounds” (**Data S1**).

**Figure 3.**
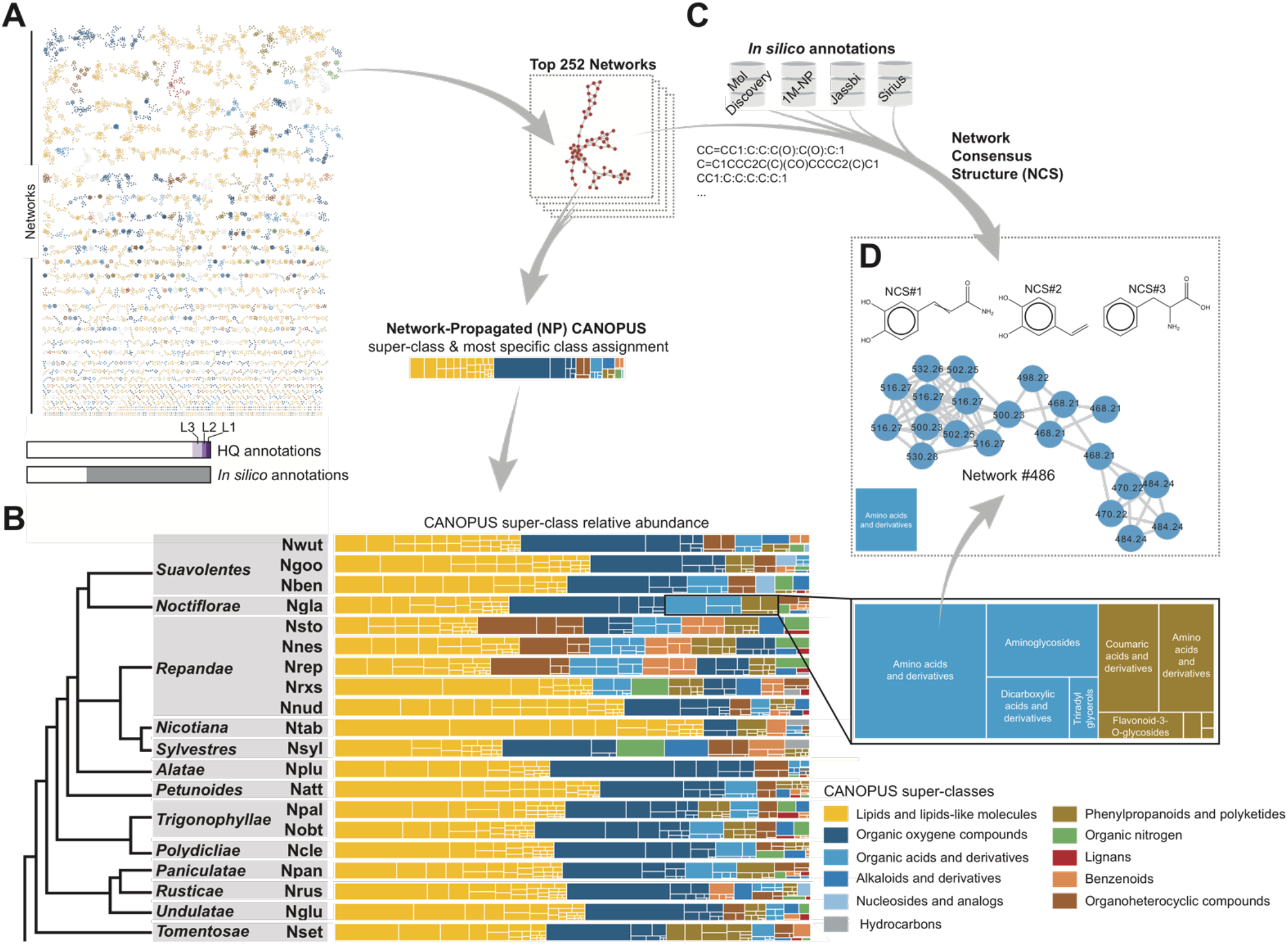
Cartography of *Nicotiana* species-level metabolic class and substructure distribution using a novel molecular network-propagated consensus substructure approach. **(A)** Molecular networking of species x tissue deconvoluted MS/MS features. The top252 molecular networks were retrieved for a minimum MS/MS pairwise cosine value of 0.7 and of 6 matching *m/z* signals. Node colors refer to network-propagated CANOPUS super-class predictions. Bars refer to the relative proportions of individual MS/MS further annotated from the three levels of annotation confidence (see **Material and Methods** section) or with databases build from *in silico* generated MS/MS spectra (see panel **C**). **(B)** Treemap visualization of species-level super-class and most specific class distribution. Colors denote for different NP-CANOPUS super-classes, with each individual uniformly colored rectangles depicting most-specific classes hierarchically classified as part of a NP-CANOPUS super-class. A close-up view on two super-classes (“Organic acids and derivatives” / “Phenylpropanoids and polyketides”) detected in *N. glauca* (Ngla) is presented. **(C)** Network Consensus Structure (NCS) computations from hits obtained from the interrogation of *in silico* generated MS/MS spectra (**Fig. S10**). Hits obtained for each MS/MS feature-level database search within a network were compiled input to compute a consensus (sub)structure for each network. **(D)** NCS computed for network #486 whose MS/MS features were classified in **(A)** as those of “Amino acids and derivatives”. The library of feature/network/NP-CANOPUS/NCS associations is reported in **Data S1, S3** and **S4**.

### Deep metabolome annotation empowered by a multi-inference approach incorporating a 1 million natural product *in silico* spectral database and consensus substructure computations

The previous analysis indicated a critical need not only for broadly increasing feature annotations beyond CANOPUS class predictions but also for gaining structural insights into core (sub)structures underlying molecular networks’ topology. As outlined in a recent review (*40*), substructure annotation provides information on functional groups, building blocks, or scaffolds within a chemical structure. This level of information is complementary to compound class prediction, most commonly addressing biosynthetic origin and/or compound physico-chemical properties. To propel substructure identification in our dataset, we first optimized a multi-inference annotation pipeline (**Fig. S2** and **S5**). Briefly, feature spectra were first queried against an in-house *Nicotiana attenuata* SM MS/MS database (NaMS, entries resulting from the analysis of purified SMs) and the GNPS library, the resulting hits being referred respectively to as annotation levels 1 to 2 according to the Metabolomics Standard Initiative nomenclature (*41*). Interrogation of these two experimental spectral databases provided hits for 4% of the MS/MS features (**Fig. 3A**). Level 3 of the annotation nomenclature regrouped class-based annotations mostly derived from manual inspection of network-level hits (5%). To circumvent limitations in the chemical space covered by these two experimental databases, spectral interrogations were conducted in parallel against *in silico*-predicted MS/MS spectral libraries using both molDiscovery which predicts MS spectra of small molecules on-the-fly and scores their probabilistic modeling (*24*), and a combination of CFM-ID and MatchMS. To further expand the power of this approach beyond the chemical space of the molDiscovery built-in library, we computed MS/MS spectra for the 429 natural products reported in a recent *Nicotiana* phytochemistry review (*32*) and, more importantly, we undertook the development of an *in silico* spectral library for about 1.1 million natural products (1M-NP).

A comprehensive description of the creation of the 1M-NP *in silico* spectral library and of its architecture is reported as **Supplementary Text** (see also **Fig. S6 and S7)**. The capacity of such *in silico* spectra-based approach to increase the annotation coverage of plant SM profiles has initially been exemplified in a pioneer study by Allard et al. (2016), but was restricted to chemical entries (∼ 220,000) retrieved from the copyrighted Dictionary of Natural Products (http://dnp.chemnetbase.com). Here, we concatenated chemical structures derived from several public natural product libraries (**Table S3**), which resulted, after filtering out duplicated InChI representations and CFM-ID-based computation of composite MS/MS spectral predictions, into 1,066,512 unique MS/MS spectra that covered a vast proportion of the natural product chemical classification proposed by NP-classifier (*43*). As CFM-ID version 4.0 computations returned slightly different MS/MS spectra for stereoisomers – see MS/MS spectra predicted (+)-/(-)-shikonin and (+)-/(-) thalidomide in **Fig. S8** –, stereoisomers were kept in the library. Altogether, this important computational delivery of this study represents, to the best of our knowledge, the largest natural product-derived *in silico* spectral library and is now available for spectral interrogation as part of the GNPS ecosystem (**Data and Material Availability**).

The above-described multi-query approach of the 17901 features from our dataset retrieved annotations for 57 % of these features, with 9 % hits for priority levels 1-to-3 (**Fig. 3A**). To maximize structural insights that could be gained from this deep annotation, we finally computed the top most common substructures (referred to as Network Consensus Structure, NCS) based on feature annotations for each of the FBMN molecular networks that did not contain any level 1-to-2 annotations. Consensus structure computational prediction relies on a new algorithmic approach that employs hits obtained from *in silico* MS/MS spectral databases (See description in the **Method section** and **Code Availability and Description**). The NCS strategy is illustrated in **Fig. 3C-D** with top NCS hits for network #486 whose MS/MS features were initially classified as “Amino acids and derivatives” by CANOPUS. A complete overview of the top NCS predictions is summarized in **Data S3**. Altogether, this unique combination of different computational approaches generated a multi-modal SM cartography that can be navigated from CANOPUS-based ontology predictions down to sets of molecular networks connected to a given class level and further down to predicted shared substructures within these networks (**Data S3 and S4**).

### Exploring the chemical substructure basis of *Nicotiana* section and species-level SM specialization

Next, we navigated the SM cartography to further dig into the inter-species chemodiversity variations that were detected from the species-level α-diversity (**Fig. 2**) and CANOPUS treemap analyses (**Fig. 3**). To rigorously infer statistical associations between species and particular CANOPUS “super-class” / ”most specific class” predictions, we employed non-metric multidimensional scaling (NMDS). NMDS is a powerful ordination technique in information visualization that is frequently employed in ecology to spatially represent interconnections among species or communities based on a series of univariate descriptors (*44*). The strength of this statistical approach is that it allows to efficiently collapse the information from multiple dimensions (here summed peak areas and connected CANOPUS predictions) into a limited number of descriptors exhibiting high-confidence statistical associations to species. Using NMDS, we computed projections of species and CANOPUS predictions as intrinsic variables and extracted strongest associations based on *P*-values < 0.05 (Permutation tests) and minimal cosine scores for angular distances between these two set of entities in NMDS projections (**Fig. 4A, Data S2**). A hierarchical clustering analysis of previously extracted most significant associations resulted into four main clusters referred to as Family Clusters (FC) (**Fig. 4B**). Distribution of these associations was not directly consistent with the species-/section phylogeny and thereby indicative of gains and losses in species/section-level capacities for the abundant production of specific SM core structures. Family Cluster 1 (FC1) regrouped predictions associated to *O*-acyl glycerol structures that appeared to be prevalent within species of the section *Suaveolentes* and to a lower extend in the *Petunoides, Polydicliae, Paniculatae* and *Rusticae*. In accordance with the pronounced expansion of this compound class in the *Nicotiana* genus (*11*), *O*-acyl sugar predictions enriched in FC2, exhibited widely distributed significant species associations throughout the genus. Such associations were remarkably absent for the section *Repandae*, with the exception of *N. nudicaulis*. Strong associations with predicted terpenoid structures caught our attention when inspecting FC3. Most distinctive ones were detected for sections *Nicotiana* and *Sylvestres* as well as for more distantly related sections *Undulatae* and *Tomentosae*. FC4 mostly captured associations with phenylpropanoid-derived substructures and alkaloids, the latter further emphasizing on the richness of alkaloid metabolism in the *Repandae* section.

**Figure 4.**
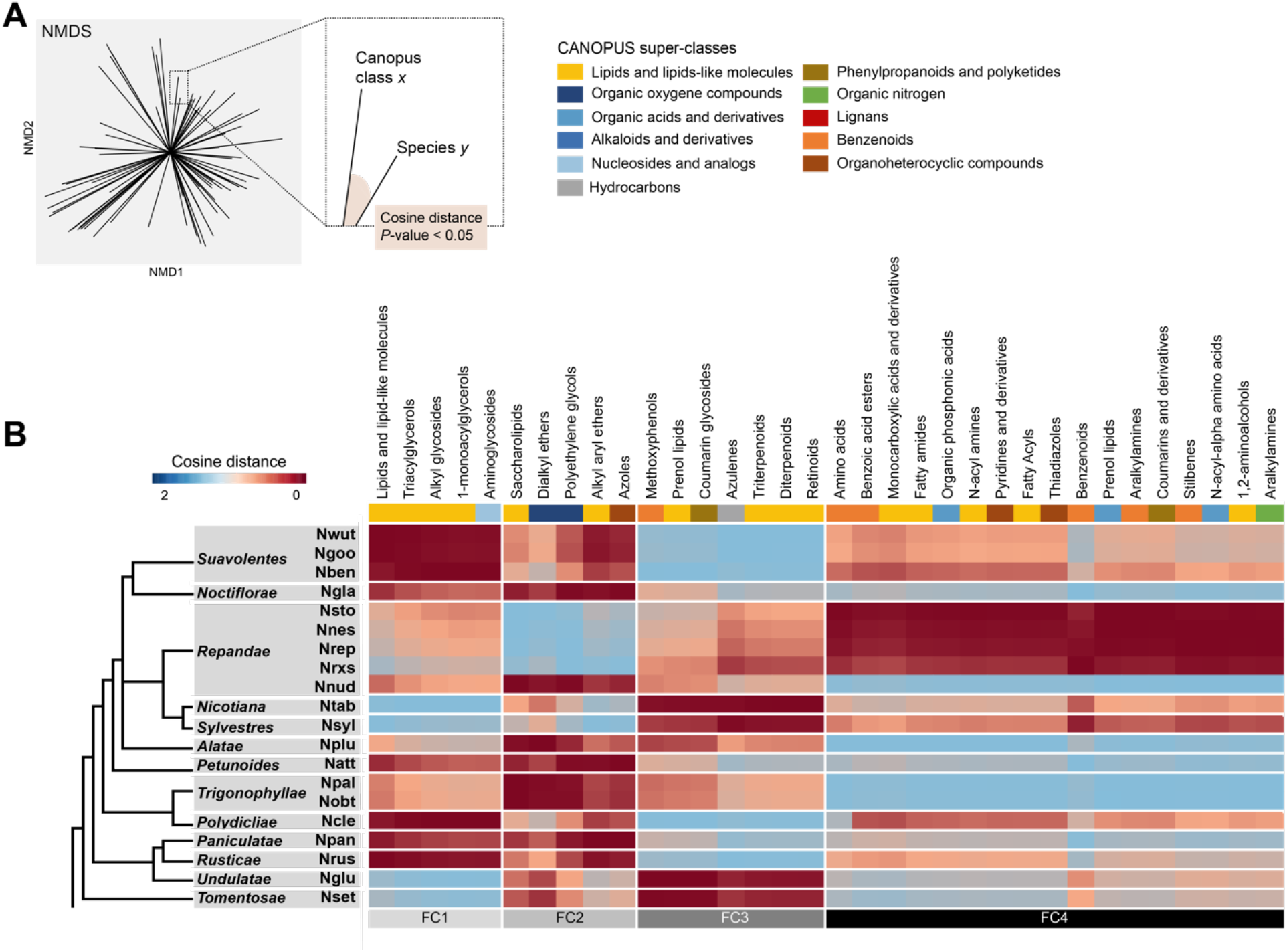
Non-metric multidimensional scaling reveals main statistical trends of *Nicotiana* section and species-level metabolic specialization. **(A)** Non-metric multidimensional scaling (NMDS) was used to infer directionalities, followed by the calculation of intrinsic variables to test for statistical significance (*P*-value [999 permutations] lower or equal to 0.05), in the association between species and CANOPUS super-class and most specific class predictions (CANOPUS, **Fig. 2**). All *P*-values and cosine distances are summarized in **Data S2. (B)** Heatmap representation (based on cosine distances) of statistically significant associations between species and NP-CANOPUS predictions for “most-specific classes” (colored according to upper-level “super-classes”). A hierarchical clustering analysis was conducted to group similarly distributed CANOPUS predictions, thereby emphasizing on four highly distinctive clusters referred to as metabolic family clusters (FC).

A detailed interpretation of these species/section metabolic specificities requires a simplified access to the underlying MS/MS fragmentation schemes. The latter can typically be approached through MS2LDA, an unsupervised method to extract common patterns of mass fragments and neutral losses, referred to as mass motifs, from collections of fragmentation spectra (*22*). From this analysis, we retained 76 mass motifs that best depicted the structural diversity within our dataset as confirmed by hierarchical clustering (resulting in clusters of co-varying mass motifs) and mapping of enriched CANOPUS predictions for each mass motifs (motif-level propagation of CANOPUS predictions) (**Fig. 5A, Fig. S9**). In analogy to the critical role of conserved domain/motif inferences in protein structure-activity studies, mass motif inference offers a dimensionality reduction perspective on recurrent fragmentation patterns derived from particular substructures. This approach is however often limited by the scarcity of structurally annotated mass motifs in MS2LDA libraries. An asset of our approach is that it mutualizes the previously described SM cartography to mine most interesting mass motifs (**Fig. 5B** and **Data S5**). For instance, we confirmed the presence in motif cluster 1 (MC1) of a mass motif (Strepsalini_110) which was characteristic of the *O*-acyl glycerols specific to *Suaveolentes*. MC1 also contained motif #631 and motif #254 characteristic of steroidal glycoalkaloids and that were strikingly specific to *N. plumbaginifolia*. Motif #646, present in the second cluster (MC2) captured the complete diversity of 17-HGL diterpene glycosides, allowing to efficiently explore tissue-specificity for this compound class. MC4 contained a motif (motif #37) with fragments indicative of hydroxycinnamic acid substructures derived from a network of *O*-phenolic glycosides. Similarly using inferences derived from these different computational approaches, we could efficiently inspect motifs corresponding to previously mentioned *N-*hydroxycinnamoyl-spermidine conjugates specific to *N. glauca* (MC5, motif #473), di- and triterpenoids abundantly found in *N. tabacum* (MC5, *e*.*g*. motif #555, #euphorbia_350) and mono-, sesqui- and diterpenes (MC5, motifs #558, #675 and #576 respectively) in sections *Nicotiana* and *Sylvestres* as well as *Undulatae* (**Fig. S10**). As previously implemented for molecular networks (**Fig. 3**), mass motifs can also be used for consensus substructure computations (Motif Consensus Structure, MCS), the latter providing a further mean to circumvent the scarcity mass motif annotation in MS2LDA libraries. All 76 MCS computations, combined with CANOPUS predictions and manual curation, are presented in **Data S6**.

**Figure 5.**
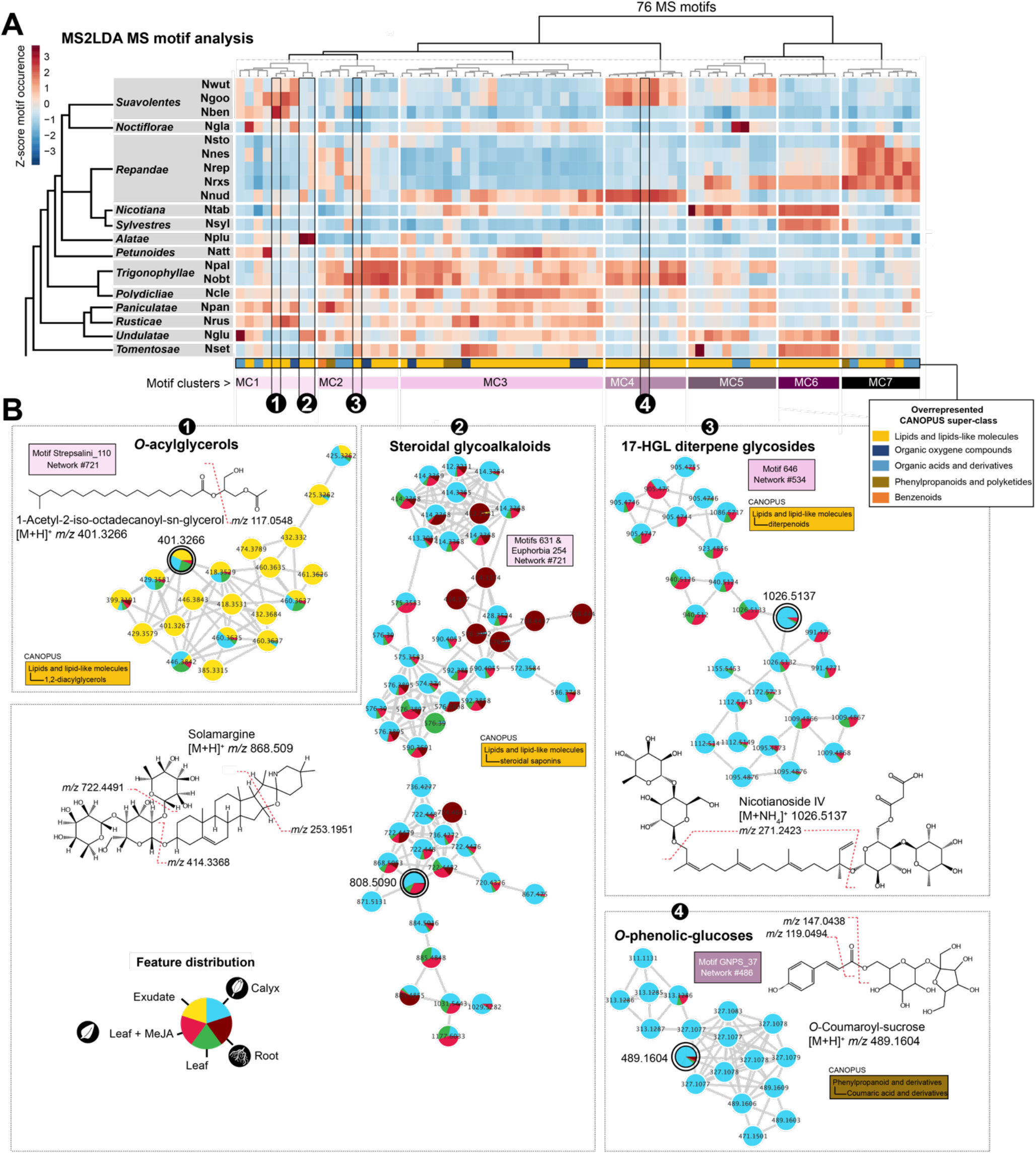
A minimal set of MS motifs captures substructure diversity in *Nicotiana* chemotypes. **(A)** Hierarchical clustering analysis (HCA) based on the species-level motif count (Z-score normalized) of top76 mass motifs inferred by unsupervised decomposition of overall MS spectra via the text-mining program MS2LDA. Species *x* tissue motif counts matrices can be explored within **Data S5**. Motifs clusters (MC) extracted from the HCA approach refer to clusters of tightly covarying MS motifs. A Principal Component (PC) analysis (2 first PCs) based on species-level MS motif relative intensity and loadings exerted on sample PC coordinates by each MS motifs, highlighted the strong resolving power for species grouping of these MCs (**Fig. S9**). **(B)** Strategy for MS motif-guided exploration of substructure enrichment in particular molecular networks. MS motifs are selected based on their peculiar species/section-level distribution, annotated using MS fragmentation curation and connected molecular network are finally visualized. Node colors denote for the species-overall feature relative abundance in the analyzed tissues. Rectangles report network and MS motif ids, their colors refer to MC. A representative high confidence predicted structure per network (connected to the double circled node) is presented with annotation of the MS motif main fragments. Additional examples are presented as part of **Fig. S10**. Overall MS motif data are reported in **Data S6** and **S7**.

### N-acylnornicotines (NANNs) as case-study for structural diversity expansion in Repandae allopolyploids

In the following, we exemplify using the case-study of NANNs, how the *Nicotiana* genus SM cartography and connected annotation resources can be exploited to gain novel (bio)chemical and evolutionary insights into specific SMs. NANNs have been described as leaf exudate allopolyploidy-mediated innovations specific to the *Repandae* section (*35*). In our data-platform, NANNs’ structural diversity was readily inferable from mass motif #433 (MC7) that included the two main nornicotine substructure molecular fragments at *m/z* 132.0825 and at *m/z* 149.1075 (**Fig. 6A**). Inspection of this motif retrieved a far greater structural diversity than previously reported, with 102 of annotated NANNs, not counting novel non-canonical NANN structures with three N (NANNs integrating an aminated fatty acyl chain) or three O atoms (di-hydroxylated NANNs) or those built on an anatabine scaffold instead of nornicotine (**Data S8**). This NANN structural diversity directly translated from variations at the fatty acyl moiety level, with the presence of *iso*-/*anteiso*-branched or straight C_1_ to C_18_ chains, with or without hydroxyl groups. As their structure had not been unambiguously identified in previous phytochemical reports (*35*), the most abundant hydroxy NANNs were purified and elucidated by NMR to confirm the unusual position of the hydroxy group at position 3 (**Fig. S11, Supplementary Text**).

**Figure 6.**
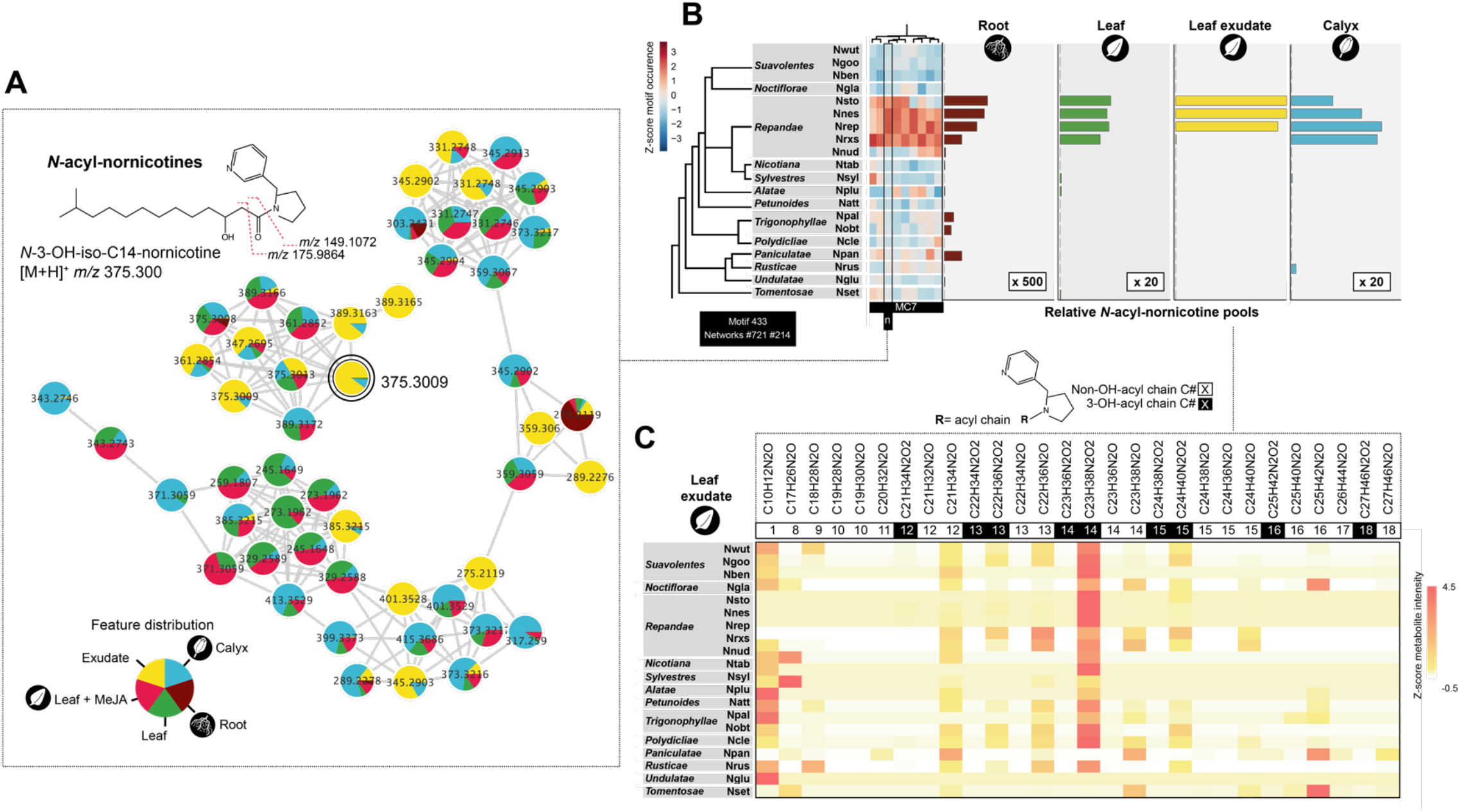
Navigating MS motifs pinpoints on a diversity of *N-*acylnornicotines that dominate leaf surfaces of *Nicotiana* section *Repandae* species. **(A)** Main molecular networks extracted connected to MS motif 433 (MC7, **Fig. 5A**) characterized by a strong relative abundance in *Repandae* species. NMR-elucidated *N-*acylnornicotine (NANN) structure (see further NMR-elucidated NANNs in **Fig. S11**), with fragment annotations captured by the NANN MS motif, for the MS/MS feature represented by the double circled node. Node colors denote for the species-overall feature relative abundance in the analyzed tissues. **(B)** Total NANN pools (relative to maximum in *N. nesophila* exudates) as inferred from MS/MS features of MS motif 433 (**Data S8**). **(C)** Species-level NANN elemental formula distribution (Z-score normalized) and indication of the acyl chain length and of its 3-hydroxylation.

Total NANN pools were extremely high in leaf exudates and in trichome-rich calyces of the *Repandae* species, but at barely detectable levels in *N. nudicaulis* (**Fig. 6B**). Most surprisingly, our data mining revealed that roots harbored a previously unexplored diversity of NANNs, albeit at almost 2 orders of magnitude lower than in leaves, and with very different chemotypes (**Fig. 6B**). In this respect, cross-tissue comparisons of fatty acyl moieties among NANN chemotypes indicated a general tendency towards shorter chain NANN (most notably C_8_-nornicotine and formyl-[C_1_]-nornicotine) accumulation in root tissues (**Fig. S12**). A closer inspection of previously noted non-canonical NANNs captured by this exploratory approach led to the formulation of structural assignments for 4 structures harboring a second intra-chain hydroxyl group, and 4 additional ones bearing a third N atom as part of an intra-chain amine group (**Fig. 7A**). These non-canonical NANNs were purified; but due to insufficient yields, their structure could not be further interpreted by NMR. In agreement with the presence of a third N prone to be positively charged, these non-canonical NANNs mainly appeared in the form of their [M+2H]^2+^ and exhibited higher polarity than regular ones. Features corresponding to these non-canonical NANNs shared with canonical ones the mass motif #433 associated with the nornicotine backbone fragmentation, but were located in different molecular networks (**Fig. 7A**) that were specific to the *Repandae* section (**Fig. S13**). These *Repandae* non-canonical NANNs were further analyzed by ultra-high resolution MALDI MS imaging experiments conducted from leaf cross-sections of *N. nesophila*. These analyses supported their uniform distribution within the leaf lamina, the corresponding MSI images overlapping with those of well-known lamina-distributed SMs such as chlorogenic acid, and not specifically on the leaf surfaces as for canonical NANNs (**Fig. 7B, Fig. S14**).

**Figure 7.**
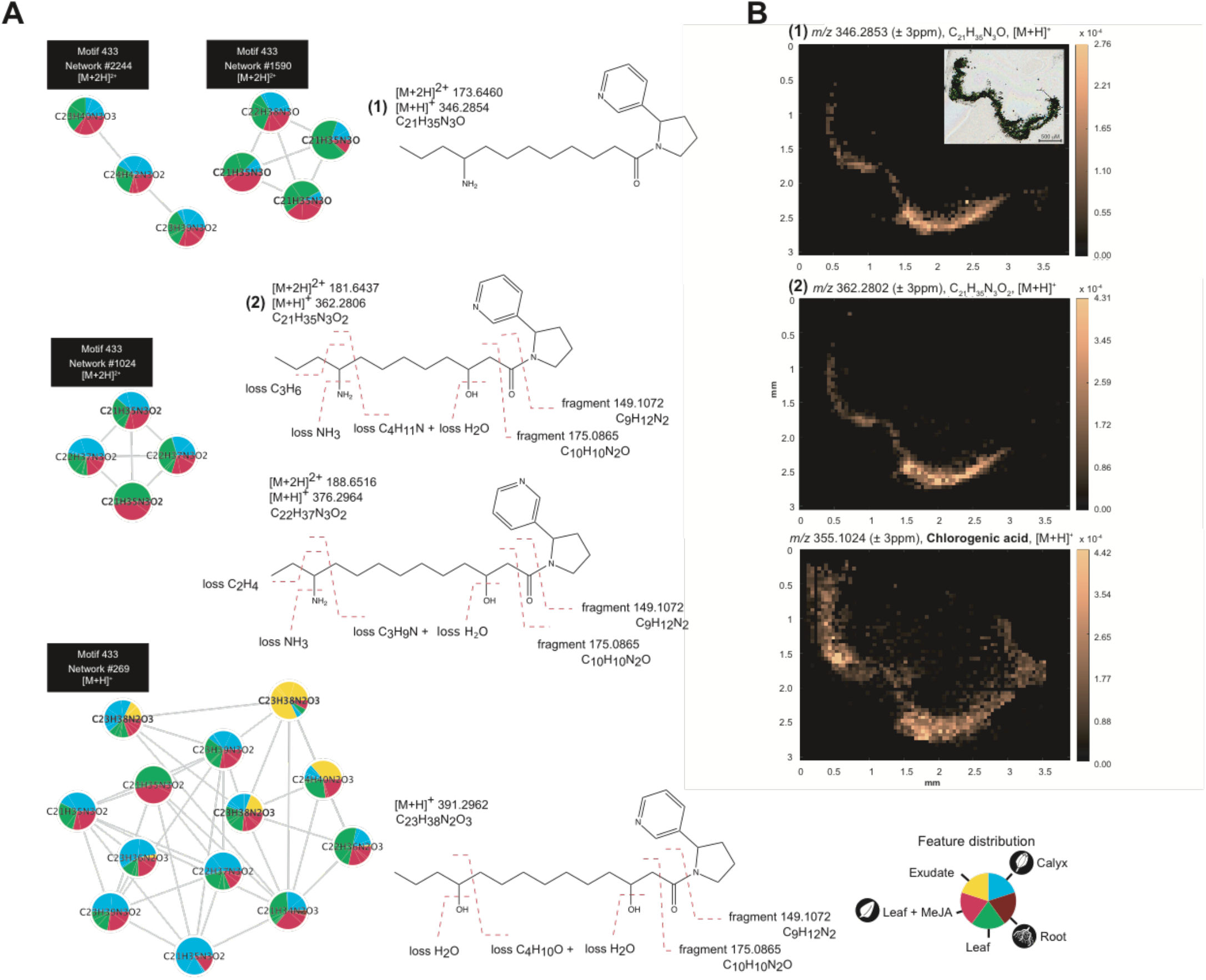
Characterization of non-canonical leaf lamina NANNs specific to the *Repandae*. **(A)** Molecular networks and fragmentation characterization of 3N-containing and di-hydroxylated NANN specific to the *Repandae* (**Fig. S13**). Node colors denote for the species-overall feature relative abundance in the analyzed tissues. **(B)** MALDI MS images depicting spatially-resolved relative abundance of selected metabolites in a leaf cross section of *N. nesophila*. Insert in the first image corresponds to the optical image of the matrix-embedded leaf cut used for MALDI MSI. The two first images correspond to the MSI data for two 3N-containing NANNs: m/z 346.2853 (± 3ppm, C_21_H_35_N_3_O, [M+H]^+^) and m/z 362.2802 (± 3ppm, C_21_H_35_N_3_O_2_, [M+H]^+^) exhibiting a quasi-uniform distribution within the complete leaf section and comparable to that of chlorogenic acid (third image, *m/z* 355.1026 ± 3ppm). Selected MSI data are presented for additional *N. nesophila* metabolites in **Fig. S14**.

### NANNs evolutionary diversification predates Repandae polyploidy formation

Our data strongly challenged the previous view that NANN biosynthetic capacity strictly arose as part of the allopolyploidy event at the base of the *Repandae* and that as such NANNs could be considered as a transgressive metabolic trait to this section. Indeed, **Fig. 6** shows that the NANNs’ diversity pervades the different *Nicotiana* sections, albeit at extremely low levels in all the species examined additionally to the *Repandae* section. Obviously, complete leaf extracts of *N. nesophila* (*H*=3.25, 53 NANNs) and *N. repanda* (*H*=3.17, 49 NANNs) exhibited the overall greatest NANN α-diversity values (**Fig. S15**). Of all leaf exudate samples examined, the NANN α-diversity calculated for hybrid *N. repanda X sylvestris* (*H*=3.03, 18 NANNs) was the highest, which reflected a balanced distribution among NANN relative intensities in this sample. By clear contrast, lowest NANN α-diversity values were detected for leaf exudates of *N. repanda* (*H*=0.42, 24 NANNs), *N. stocktonii* (*H*=0.39, 22 NANNs) and *N. nesophila* (*H*=0.54, 24 NANNs), which further indicated, besides the high NANN biosynthetic capacity in these species, their exacerbated specialization towards C_14_-OH-nornicotine exudation. In this respect, while the NANN chemotypes of the leaf exudates of almost all of the focal species were characterized by the dominance of this particular NANN, *N. rustica* and *N. setchellii* were noticeable exceptions, being dominated by C_16_-nornicotine (**Fig. 6C**) and *N. glutinosa* for its exclusive accumulation of formyl-nornicotine. As previously noted (**Fig. 6B**), roots of almost all species harbored a rich diversity of NANN, particularly exacerbated in *N obtusifolia* (*H*=2.26, 8 NANNs), predicted as one of the closest diploid progenitors to the *Repandae* section. Altogether, a most parsimonious explanation to the evolution of the NANN pathway was that it predates *Repandae* formation. Such an evolutionary scenario appeared to be supported in all tissue-level ancestral state reconstruction (ASR) analyses carried out based on a *matK*-based species tree and with total NANN levels expressed as discrete states (**Fig. 8**). The ASR analysis computed from total root NANNs in combination with tissue-level NANN chemotypes, further suggested that the last common ancestor to the examined species had a consequent root-based NANN accumulation capacity.

**Figure 8.**
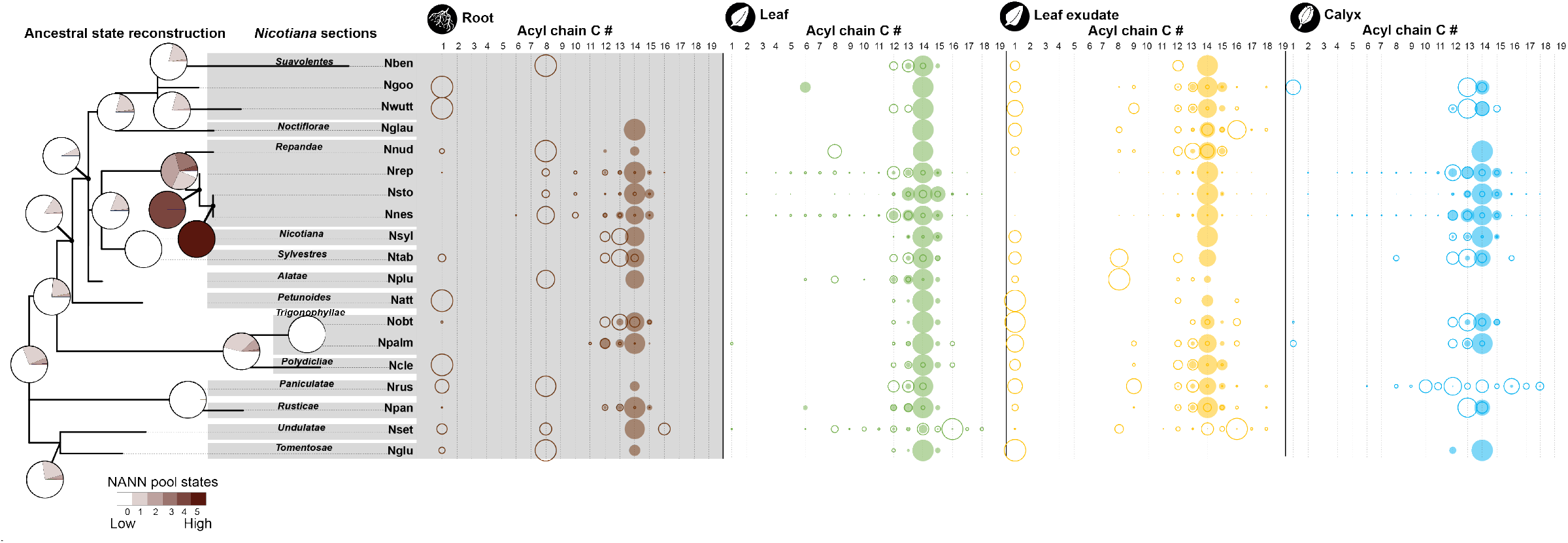
Ancestral state reconstruction and structural diversity analysis indicate that NANNs predate *Repandae* speciation and a major root-to-shoot compositional shift. Total root NANN pools of the focal species were transposed as relative scaling into an ordered trait (total states colored from white to dark brown) and used as input for ancestral state reconstruction using the MBASR software with default settings. The species tree was constructed from *matK* as described in (*71*). Bubble plots on the right part of the figure depicts relative NANN fatty acyl chain distribution with indication of fatty acyl chain carbon number (**Fig. S12**), for total NANN pools see **Fig. 6B**. Bubble size denote for relative acyl chain level within the NANN pool of a species and *per* tissue. Color-filled bubbles refer to hydroxylated NANNs.

## Discussion

Lineage-specific reconfigurations in rapidly evolving sectors of a plant specialized metabolism can be transparent at the genomics/transcriptomics levels for which most evolutionary studies on adaptative traits are conducted. This stresses the obvious fact that the power of genomics-driven evolutionary inferences on plant SM pathways critically relies on the chemical classification of metabolites part of these metabolic sectors as well as on the phylogenetics contextualization of this information. To tackle this issue, the open-source computational metabolomics approaches presented here are propelled by a broadly transposable multi-inference annotation that maximizes the coverage of substructure predictions, thereby resulting into an unprecedented cartography of SM diversity in the *Nicotiana* genus linking species-level SM prevalence to particular substructures. With this workflow, we notably shed light on the structural diversity and phylogenetics distribution of NANNs, a gain-of-function defensive innovation previously thought to have evolved with *Repandae* allopolyploids speciation (*38*).

A major challenge in MS metabolomics remains to reach broad structural annotation (“deep metabolome annotation”) and substructure discovery beyond chemical class predictions and the dereplication of previously identified SMs, which is the most frequent outcome of molecular networking-based data exploration. In particular, with the use of heterogeneous computational annotation tools and that of querying highly diverse experimental and *in silico* MS/MS database comes the inherent difficulty of systemically prioritizing and/or merging the minimal set of most reliable annotations collected from these inferences. MolNetEnhancer has been developed to more efficiently combine outputs from molecular networking, MS2LDA as well as *in silico* and chemical classification tools (*45*). However, substructure discovery from MolNetEnhancer outputs is strongly hampered by the scarcity of annotated motifs in the Mass2Motifs database embedded into MS2LDA, many of which additionally translating from relatively unspecific fragmentations (e.g. water, methyl, hexose losses). Only 24 of the 76 mass motifs retained for further analysis had partial annotation hints in the MS2LDA Mass2Motifs database (**Data S5**). To significantly improve substructure discovery and annotation, we implemented two complementary approaches. On the one hand, we propagated CANOPUS predictions at mass motif level (MP-CANOPUS) by computing frequencies in “super-class/sub-class/most specific class”, and combined this information with mass motif co-regulation analysis (**Fig. 5**). The second approach implemented for substructure analysis involved advanced maximum common substructure calculations to integrate annotations from multiple tools on a network (NCS) or motif level (MCS). Overall, we obtained 349 NCS or 303 MCS predictions for the whole data-set (**Data S3 and S6**). Maximum common substructure computation for substructure prediction had been employed in one of our previous studies to cluster candidate structures obtained by the MetFrag searches among co-regulated herbivory-induced metabolites (*46*) and is also one of the processing steps within the Network Annotation Propagation tool of the GNPS web-platform (*47*). Altogether, we advocate that the NCS/MCS approach implemented here has three main advantages: (*i*) it is an efficient mean of summarizing common substructure within the diversity of outputs from database queries as SMILES strings, (*ii*) it can used as input to reveal substructures statistically associated with intense chemodiversification in a given species, and (*iii*) it provides structural guidance during the manual interpretation of mass motifs or molecular network. In this respect, our study led to the curation of 76 mass motifs (**Data S5**). Such effort is important to empower supervised search of mass motifs which is already possible in MS2LDA and which will be greatly facilitated with the recent release of the MS2QUERY tool (*48*).

A very important delivery of our work is the development and public sharing of the 1M-DB which is, to the best of our knowledge, the largest *in silico* spectral database. This approach resulted into 5-fold more hits (annotation of 57% of the total features), than experimental spectral database interrogation alone. Data of the 1M-DB can currently be accessed and interrogated from the GNPS platform. The size of this data-set can represent a challenge for MatchMS-based queries, which can nonetheless be locally implemented with reasonable computing capacity with the parallelized script (**Code Availability and Description**) provided with our study. It is therefore foreseeable that the efficiency of the interrogation of the 1M-DB will strongly benefit from up-to-date optimization of MatchMS parallelization as part of future version releases. Multiple tools have been developed in recent years to produce hypothetical MS/MS spectra (*23–26, 49*). A more recent development in this area is that of QCxMS which provides, in our experience, very high-quality spectra. This program is currently too much computationally demanding and could not be transposed to the scale of this study, besides the computation of MS/MS predictions for the 429 structures of the Jassbi database and using a limited number of fragmentation trajectories (Zenodo link, https://doi.org/10.5281/zenodo.6536010). One promising direction for improving the confidence of such *in silico* fragmentation-based annotation is exemplified by the recently developed COSMIC workflow that incorporates a confidence score consisting of kernel density *P-*value estimation from a decoy library and a support vector machine algorithm (*50*). With the increasing quality of MS/MS predictions, one interesting perspective could be to extract mass motifs from them and thus directly infer fragment substructures produced from known structure *in silico* decomposition.

In terms of structural information, rhe SM metabolic cartography generated in this study goes far beyond to a recently published chemotypic classification of the *Nicotiana* genus which mostly consisted in the dereplication of primary metabolites such as steroids and only a few SMs (*51*). In our opinion, this data platform and our SM cartography provide complementary views on the metabolic diversity of this genus. Noteworthy, the aforementioned study did solely focus on leaf metabolomes, while ours and previous studies (*8*) unambiguously indicated the importance of “screening” multiple tissues to capture a broader SM diversity picture. In this respect, we demonstrated that expanding the analysis at the multi-tissue level (by combing tissue-level molecular network information) resulted into a “phylometabolomics” tree that captured shared SM biosynthetic potential among closely-related species with more resolution (**Fig. 2**). Beyond simple presence/absence of SM classes which has been a traditional focus of chemotaxonomic studies, the fact that structural diversity can nowadays be more efficiently accessed with computational MS metabolomics opens novel research avenues for understanding the evolution of SM, as implemented in a recent survey of the SM synapomorphies and homoplasies in the Malpighiaceae family (*52*). Information theory Shannon statistics transposed to MS feature analysis or individual metabolites can also provide an efficient means of contrasting metabolic diversity among the metabolic profiles to examine evolutionary ecology theories and contextualize those at relevant taxonomic scales (*53*). By employing α-diversity analysis, we confirmed that roots exhibit, under our analytical conditions, the most specific metabolomes, a pattern which had been previously detected in a study focusing on *N. attenuata* as the sole model species (*8*). α-Diversity scores further varied in-between species, thereby indicating variations in constitutive SM biosynthetic capacities and/or constitutive vs stress-induced investments into SM production. In this respect, we further observed that these inter-species variations in MeJA inducibility were negatively correlated with α-diversity scores constitutive leaf metabolome. This trend is reminiscent of the inter-species patterns detected from the comparative analysis of early herbivory-induced transcriptomes for 6 *Nicotiana* species (*54*), and may reflect physiological trade-offs between constitutive vs inducible metabolic diversity maintenance.

Many interesting novel biochemical insights worth to be pursuing by gene function studies, were extracted from the SM cartography produced from this study. Our analysis notably detected the presence of mono-*O*-acylglycerols (classified under the CANOPUS most specific class 1-monoacylglycerols) specifically on the leaf surfaces of the section *Suaveolentes* and at lower abundances in the *Rusticae*. Besides its well-known housekeeping function in the synthesis of di- and tri-*O*-acylglycerols via the action of GPAT enzymes (*55*), the latter compound class has been poorly investigated regarding its presence on plant aerial surfaces. Main reports on the possible defense-related functions of this compound class derive from studies on their presence as abundant surface metabolites on the calyx of several Scrophulariaceae species (*56*), and from a unique report for the *Nicotiana* genus describing these compounds as efficient chemical glues against small insects on the leaf surfaces of *N. benthamiana* (*57*). The prevalence of this compound class in the *Suaveolentes* section, in particular in *N. benthamiana*, along with the here-described high levels of *O*-acyl glucoses (*58*) could point to an interesting case-study to functionally examine the biochemistry and evolution of this pathway and compare it with that of the thoroughly investigated and structurally reminiscent *O-*acyl sugars (*11*). Our analysis also revealed subtle tissue-level chemotypic variations within *O-*acyl sugars networks. Apart fro, confirming previously detected strong cross-species variations in structural diversity, inspections of these networks also pinpointed that some of these *O-*acylsugars are present at low levels in roots (**Fig. S10**). This could further illuminate recent work on the predicted role of these SMs in plant-soil microbiome interactions (*15*). Our SM cartography also provided a far greater species *x* tissue resolution on terpene-related classes’ distribution compared to tendencies previously sketched in *Nicotiana* studies that targeted trichome-based cembrene diterpene (*59*) and 17-HGL-DTG (*6*). Our study revealed for these two classes of diterpenes, pronounced expansions of structural diversity and significant associations with the *Nicotiana, Sylvestres, Undulatae, Tomentosae, Trigonophyllae* (17-HGL-DTG) sections that include species in which emblematic structures of these compound classes had been originally detected (*6*). An unexpected result was the detection, at large levels in *N. plumbaginifolia* and to a minor extent in *N. glutinosa*, of steroidal glycoalkaloids, emblematic of the *Solanum* genus and whose presence is considered as erratic in other Solanaceae genera. Within the structurally rich network of steroidal glycoalkaloids identified in our study, the dereplication of solaplumbin *m/z* 722.4479, ([M+H]^+^, C_39_H_64_NO_11_) is supported by old phytochemistry reports (*60*). Such unexplored patchy distribution of steroidal glycoalkaloids within the Solanaceae provides exciting foundations for future evolutionary biochemistry studies.

The fact that α-diversity scores, independently of the tissue type, were not reflecting direct metabolic additivity in allotetraploid species. This, along with unique metabolic characteristics as compared to their closest extent diploid progenitors, could be reminiscent of patterns observed when inspecting complex reconfigurations of floral morphological and associated metabolic traits in *Nicotiana* allotretraploids (*34*). Due to their previously reported absence in *Repandae* closest diploid progenitors (*Nicotiana sylvestris* and *Nicotiana obtusifolia*), NANNs have often been considered as “transgressive” metabolic traits derived from the *Repandae* allopolyploidization. In our study, we annotated 102 NANNs, including 6 first elucidation by NMR, and discovered NANN-related structures built from anatabine as a backbone, and the presence of novel NANNs leaf lamina-based restricted to *Repandae* and incorporated uncommon aminated fatty acyl moieties. Above all, our study indicates that the NANN biosynthetic capacity predates the *Repandae* section formation. However, a main innovation of *Repandae* species is their capacity to accumulate very high level of canonical NANNs on their surfaces as well as N_3_-containing NANNs in their leaf laminas. These data provide rigorous support to old literature that reports anecdotal evidence (*61, 62*) for low amounts of short (-formyl, -acetyl) and middle (C_4_-C_8_) chain length NANNs present in other *Nicotiana* species (*63*). Interestingly, *N. obtusifolia*, considered as a closest extant female progenitor to *Repandae*, is one of the *Nicotiana* species that accumulates the largest nornicotine-to-nicotine ratio in its leaves (*64*). Another interesting observation to pursue is that *N. sylvestris*, the closest extant male progenitor to *Repandae*, is thought to have contributed to several allopolyploidization events in the genus *Nicotiana*, many of which being able to accumulate greater NANN amounts than the other species tested in this study. As such, our data suggest a more complex than previously thought evolution of the NANN pathway. A direct perspective will be the identification of the canonical NANN biosynthetic *N-* acyltransferase(s) which is predicted to be abundant in *Repandae* trichomes from our data and from previous phytochemical analyses on crude trichome fractions (*35–37, 65*). Our tissue cartography finally revealed a largely unexplored repertoire of NANNs in the roots of all examined species. These data and ASR analyses are in favor of shorter chain NANN production in roots being a most ancestral trait in this metabolic class. In the context of future biochemical investigations, the latter interpretation would be consistent with the fact that the accumulation of canonical NANNs onto aerial surfaces involves trichome-based *N*-acyltransferase enzymes with greater affinity for long chain fatty acyl-CoA as compared to those present in roots.

In conclusion, the fully open data and broad range of data integration approaches and provided here present an unprecedented resource to revive SM analysis in the *Nicotiana* genus and contribute to the establishment of phylometabolomics as an instrumental bottom-up approach to guide future evolutionary biochemistry studies.

## Material and Methods

### Plant material, growth conditions and treatment

*Nicotiana* species with their origin and associated accession numbers are summarized in **Table S1**. Seeds of all *Nicotiana* species were directly germinated on soil, with the exception of *N. attenuata*, for which smoke-induced seed germination was established as described previously (Krügel et al., 2002). For all species, glasshouse growth conditions were as described previously (Krügel et al., 2002). Six-to-eight weeks old elongated plants were used for all metabolomics analyses. In order to analyze the regulatory function of jasmonate signaling on metabolomics-inferred specialized metabolism classes, petioles of 2 elongated plants were treated with either 20 µL lanolin paste containing 150 µg methyl jasmonate (Lan + MeJA) or with 20 µL pure lanolin (Lan) according to Heiling *et al*. (2021). Leaf samples were harvested 72 h after treatment, flash-frozen in liquid nitrogen, and stored at -80°C until use.

### Metabolite extraction procedures for UPLC-QTOF MS

Leaf, root and calyx metabolites were extracted for UPLC-QTOF MS analysis as previously described (Heiling et al., 2017). Briefly, for leaf samples, 12 discs *per* plant (∼ 100 mg fresh-weight tissue) were flash-frozen in liquid nitrogen immediately after harvest and stored at -80°C until use. The latter frozen leaf samples were ground in a Tissue Lyzer II for 3 min at 30 Hz and metabolites extracted by addition of 1 mL of 80 % methanol, 1 h of shaking at 1000 rpm at 4°C and further kept with a gentle agitation overnight at 4°C. Samples were finally centrifuged for 10 min at 14000 g and the resulting supernatants transferred into glass vials. Root samples referred to the complete root system of about-8 weeks old plants. After soil removal, roots were rinsed in water, gently dried with paper towels and flash-frozen in liquid nitrogen. Root samples were homogenized in a Tissue Lyzer II for 4 min at 30 Hz. Metabolite extraction was conducted as above described from 200 to 400 mg root material (primary, secondary and tertiary roots). Flower calyces were collected from about 8 weeks old plants and processed for metabolite extraction using above leaf metabolite extraction conditions. To obtain leaf exudates enriched into semi-polar to apolar surface metabolites, fully elongated leaves were briefly rinsed with acetonitrile. These exudates were filtered on filter paper and completely dried under reduced pressure. Dried residues were then re-dissolved in methanol and total metabolite concentration was adjusted to 1 mg.mL^-1^, except for *Nicotiana repanda, N. stocktonii* and *N. nesophila* exudates which were diluted to 0.001 mg.mL^-1^ and 0.1 mg.mL^-1^ (see **Table S1**) in order to avoid detector saturation, due to the high levels of NANNs in these samples. Peaks areas were corrected by corresponding dilution factors.

### UPLC-QTOF MS chromatographic conditions

Methanolic extracts were analyzed using ultra-high pressure liquid chromatography coupled to high-resolution mass spectrometry on an UltiMate 3000 system (Thermo) coupled to an Impact II (Bruker) quadrupole time-of-flight (QTOF) spectrometer. Chromatographic separation was performed on an Acquity UPLC ® BEH C18 column (2.1×100mm, 1.7µm, Waters) equipped with an Acquity UPLC ® BEH C18 pre-column (2.1×5mm, 1.7µm, Waters) and using a gradient of solvents A (water, 0.1% acetonitrile, 0.05% formic acid) and B (acetonitrile, 0.05% formic acid). Chromatography was carried out at 35°C with a flux of 0.4 mL.min^-1^, starting with 10% B for 3 min, and reaching successively 20% B at 12 min, 35% B at 17 min, 40% B at 23 min, 45% B at 25 min, 50% B at 30 min, and 95% B at 40 min, holding 95% for 5 min and coming back to the initial condition of 10 % B in 3 min. These chromatographic conditions (total running time of 48 min) were previously optimized for the comparative metabolomics of methanolic extracts of Solanaceae species in one of our previous studies (Heiling et al., 2016). Samples were kept at 4°C during the sequence of injections and 5µL per sample were injected in full-loop mode with a washing step after sample injection involving 150µL of the wash solution (water:methanol, 80:20, v:v).

### Conditions for DDA MS/MS data collection during UPLC-QTOF MS analysis

The Impact II QTOF instrument was equipped with an electrospray ionization source and operated in positive ionization mode on a 50-to-1500 Da mass range with a spectra rate of 5 Hz and by further using the AutoMS/MS fragmentation mode. The end plate offset was set at 500 V, capillary voltage at 4500 V, nebulizer at 2 Bar, dry gas at 10 L.min^-1^ and dry temperature at 200°C. The transfer time was set at 60-70 µs and MS/MS collision energy at 80-120% with a timing of 50-50% for both parameters. The MS/MS cycle time was set to 2 seconds, absolute threshold to 31 cts and active exclusion was used with an exclusion threshold at 3 spectra, release after 1 min and an ion was reconsidered as precursor for the fragmentation if the ratio current intensity/previous intensity was higher than 5. MS/MS collision energy was set according to the mass from 25 V for a mass of 100 Da to 50V for a mass of 1 500 Da. The MS/MS spectra acquisition rate was further optimized, from 3 Hz to 7 Hz, according to the intensity of the observed mass. A calibration segment was included at the beginning of the runs allowing the injection of a calibration solution from 0.05 to 0.25 min. The calibration solution used was a fresh mix of 50 mL isopropanol:water (50:50, v:v), 500 µL NaOH 1M, 75 µL acetic acid and 25 µL formic acid. The spectrometer was calibrated on the [M+H]^+^ form of reference ions (57 masses from *m/z* 22.9892 to *m/z* 990.9196) in high precision calibration mode with a standard deviation below 1 ppm before injections, and re-calibration of each raw data was performed after injection using the calibration segment.

### Ultra-high resolution MS imaging data acquisition and processing

Freshly collected rosette leaves of *N. nesophila* were embedded into M-1 embedding matrix (Thermo Scientific) and frozen before cutting. Cuts were done on a transverse plane at 25µm thickness and -15°C using a cryotome FSE. Sections were deposited on indium-tin-oxide coated slides and sprayed with a-cyano-4-hydroxycinnamic acid (HCCA) matrix at 10mg/mL in 70% ACN, 0.1% trifluoroacetic acid using the HTX M5 sprayer. Nozzle temperature was set at 75°C, flow rate at 0.120mL/min, velocity at 1200 mm/min, pressure at 10 psi, gas flow rate at 3 L/min and nozzle height at 40mm. Four passes were applied with a track spacing of 3mm and a HH pattern.

Samples were analyzed with a Burker SolariX 7T Fourier transform ion cyclotron mass spectrometer at resolving power R=120,000 at *m/z* = 400. Acquisition was performed in positive ion mode on a 100-500 *m/z* mass range, with an accumulation of 0.020 s, the transfer optics time of flight set at 0.600 ms, frequency at 6 Hz and RF amplitude at 350 Vpp. The MALDI plate offset was set at 100 V, deflector plate at 200 V, laser power was set at 20%, laser shots at 100 and frequency at 1000 Hz with a small laser focus. The instrument was calibrated by multipoint correction using the peaks of the HCCA matrix (*m/z* = 379.0924, 399.0377, 401.0744, 417.0483). The regions of interest were determined in FlexImaging with a raster width of 50µm. Images of the ions of interest +/- 3 ppm were displayed in MSiReader v1.03 (*66*). The data was submitted to metaspace and is available at https://metaspace2020.eu/project/nicotiana_msi-2022

### Feature-based molecular networking of UPLC-QTOF MS data

Raw data were converted to the .mzML format using MSConvert (Version 3.0.21112-b41ef0ad4, Chambers et al., 2012). The resulting data files were then processed with the Batch Mode (See Code availability, Script **S10**) of MZMine 2.53 (*68*) and exported for Feature-based molecular networking (FBMN) analysis in the GNPS environment (Nothias et al., 2020; Wang et al., 2016) and for spectral analyses in Sirius (*21*). The resulting .mgf and .csv files were further filtered to exclude redundant none-biologically informative MS/MS features using newly developed Python Scripts **S11** and **S12** (see Code availability). The m/z signals that appear more than 5 times (± 3ppm) with a retention time coefficient of variation greater then 10 % were discarded. This filtering step excluded 11580 features (out of a total of 29481 retrieved from the MZMine-based processing), a vast majority of those corresponded to redundant features detected at high-level in solvent blanks. Finally, FBMN was performed using the modified cosine as spectral similarity metric and with standard settings (Version release_28.2, except lower precursor and fragment tolerance of 0.005 Da). Output of the FBMN analysis is available on GNPS at the following link: https://gnps.ucsd.edu/ProteoSAFe/status.jsp?task=cf822b6c7c914206941bb0b6007e7eb0

### MS/MS elemental formula and compound class predictions with Sirius

Sirius (Version 4.8.2) was used to predict elemental formulas for MS/MS precursors as well as for the deep neural network-based compound class prediction as part of the CANOPUS pipeline (*20*). Sirius commands are summarized as part of Script **S13** (see Code availability). Elemental formulas by Sirius were further processed with Scripts **S14** and **S15** (see Code availability) to restore Feature IDs and calculate the degree of unsaturation of these formulas. A main strength of CANOPUS-based class prediction is that does not involve the interrogations of spectral libraries with fragmentation spectra, thereby allowing class prediction of MS/MS features for which no database hit is retrieved and circumventing the possible issue of error propagation when false class prediction is obtained by FBMN network-level propagation from feature-derived database hits. MS/MS feature-level ontologies were retrieved from CANOPUS predictions as well as FBMN network-propagated superclass, subclass and most specific class ontologies. The latter ontology propagation was implemented using Script **S19**.

### Mass motif inference by MS2LDA

Mass motifs were inferred using standard settings of MS2LDA (Version release_23.1, Wandy et al., 2018), submitted trough the GNPS workflow. Available at: https://gnps.ucsd.edu/ProteoSAFe/status.jsp?task=6f325f462e1145bfb465c679c2ee17d6 A total of 609 motifs were assigned including already existing motifs from motifdb. To explore mass motifs assignments on a species level, a binary mass motif matrix for all tissues was created by setting features above peak area of 10.000 to the value of 1 and those below to 0 (Script **S17**). The resulting matrix was combined and presence was summed *per* tissue and then set again into a binary matrix. Following this binary transposition of motif distributions, feature presence *per* motif was determined *per* species, resulting in a motif count table (Script **S18**). This set of mass motif counts (after filtering and manual curation 76 motifs), was then normalized by motif id and clustered by hierarchical clustering using the Ward clustering method implemented in MetaboAnalyst (*69*).

### MS/MS annotation based on spectral database interrogations

We implemented a 3-pronged approach to annotate MS/MS from the interrogation of experimental and *in silico* fragmentation databases, similar as proposed in Sumner et al., 2007. Level 1 in our priority assignment of annotations corresponded to hits retrieved from experimental spectral databases and or NMR structural confirmation. Highest priority within level 1 of annotated spectra (level 1a) was given to hits confirmed by NMR in this work. Level 1b annotations correspond to hits from spectral alignments (and correspondence of precursor *m/z* values) using a local MatchMS (score above 0.65 and more than 6 matching peaks) implementation (Script **S8**, see Code availability) with the modified cosine score, from an in house high-resolution experimental MS/MS spectra database of *Nicotiana attenuata* specialized metabolites and/or manual inspection of spectra. Level 2 corresponded in our annotation approach to hits retrieved, with the cosine score from high-resolution MS/MS spectra of the GNPS database. Level 3 annotations were considered for hits from alignments with *in silico* MS/MS spectra or in the case of network propagation of hits from the experimental databases, both after manual inspection. Jobs for the recently developed molDiscovery approach (Version 1.0.0, Cao et al., 2021) were submitted through GNPS with both the molDiscovery built-in library and the Jassbi compound database created as part of this study. The Jassbi compound database (429 structures) was compiled from structures extracted from a recent *Nicotiana* phytochemistry review (*32*). *In silico* MS/MS spectra for the Jassbi compound database were also produced with the fragmentation tool CFM-predict 4.0 (*23*) (Script **S5**, see **Code availability**) database searching was performed with MatchMS (*70*) (Script **S9**).

### Consensus substructure and molecular network chemical classes

We implemented a new algorithmic approach to deal with the high number of annotations retrieved from the various *in silico* MS/MS spectral databases. To this end, we used annotations retrieved form Sirius (confidence score above 0.65), 1M-DB searched with modified cosine (score above 0.5 and 5 matching peaks), CFM-DB 1M searched with spec2vec (score above 0.5), Jassbi-CFM (score above 0.5 and 5 matching peaks) and Jassbi-molDiscovery. These annotations were retrieved at the molecular network or at the MS2LDA mass motif level in order to calculate consensus substructures for a given network (NCS) or mass motif (MCS). Main steps involved in consensus substructure calculations involved the following commands (Script **S16, S19**, see **Code Availability and Description**): (*1) fragment structures, (2) get the most common fragments, (3) select the top 50 and only keep the ones with more than 12 atoms, (4) cluster by structural similarity, (5) sort by cluster size, (6) calculate the maximum common substructure within the cluster, (7) retrieve the top 4 results*.

To harness the vast amount of structural information classified by molecular networking, we selected the top 252 networks sorted by only picking networks containing more than 10 nodes. The peak areas within these networks were summed with Script **S24**. Peak areas were normalized (Excel’s STANDARDIZE function) by cluster id and the maximum on tissue level per species was kept. The propagated CANOPUS classes were grouped their peak areas summed (Script **S28**) and the resulting data was used to create per species treemaps in Excel. A summary of the Top252 molecular networks, their calculated consensus substructures and their propagated CANOPUS classes can be found in **Data S3**. Additionally, **Data S4 and S7** allow to navigate this multi-level information at mass motif and network levels.

### Computing MS/MS-informed phylometabolomics species trees

To create MS/MS similarity-based species, referred to in the text as phylometabolomic trees, we used the data compiled as mentioned above (Script **S24**) (**Fig. 2A**) or the data from the motif count (Script **S18**) (**Fig. S9**) in order to calculate the Euclidean pairwise distances between species’ metabolomes (Script **S20**). The resulting matrix was then used to plot trees in R with the APE package using the Neighbor-Joining algorithm and bootstrapping 999 with iterations. (Script **S21**)

### Ancestral state reconstruction for the relative occurrence of N-acylnornicotines

We adapted the concept of ancestral state reconstruction (ASR) classically employed for the evolutionary analysis of quantitative phenotypic traits for the exploration of NANNs’ relative occurrence. To this end, we first constructed a phylogenetic tree of the focal *Nicotiana* species based sequences of the *matK* gene obtained from a previous study (*71*), the sequence of *N. maritma* was used to account for *N. wuttkei* position within the species tree due to unavailable genome data for the latter species. Laskowska and Berbec (2003) previously suggested the very close relationship between the latter two species as well as reported their successful hybridization in the wild. *Nicotiana setchelli matK* gene sequence was obtained from the assembly of transcriptomics data publicly available for NCBI SRA accession SRR2106530. The species tree was constructed using NGPhylogeny.fr (*73*) with default one click options and the PhyML Maximum Likelihood method. For ASR, feature intensities accounting for the species and tissue-wide NANN diversity were retrieved using the above-described mass motif characterization approach. ASR was performed with the MBASR package (Heritage, 2021; Script **S25**) on peak areas of the root that have been transformed into a ordered trait of 5 categories (**Fig. 8, Fig. S1**).

### α-Diversity analysis and CANOPUS class distance computation

The alpha-diversity was calculated for each species based on Shannon Entropy (Script **S29**) using the scikit-bio package and sample features as OTUs. The top 252 networks as mentioned previously were selected their raw peak areas summed based on propagated CANOPUS classes (Script **S28**) and then converted to integers, networks without class annotations were discarded. The vegan package was used to perform non-metric multidimensional scaling (NMDS) followed by the calculation of intrinsic variables (CANOPUS classes) with 999 permutations (Script **S30**). The resulting vectors were used to calculate the per species cosine distances (Script **S31**).

### Code availability

All scripts used in this study are available at the Github repository: https://github.com/volvox292/Nicotiana_metabolomics

## Acknowledgments

The authors would like to acknowledge the High Performance Computing Center of the University of Strasbourg for supporting this work by providing scientific support and access to computing resources. Some of these computing resources were funded by the Equipex Equip@Meso project (Programme Investissements d’Avenir) and the CPER Alsacalcul/Big Data. The authors thank N. Navrot and I. Grubor for comments on the manuscript, L. Malherbe for help with plant sample collection, J. Zumsteg for support with preparative HPLC purifications, the Plant Imaging Mass Spectrometry platform of the IBMP for instrument access, P. Dorrestein, M. Wang and M. Panitchpakdi for help with the upload of the *in-silico* 1M-DB on the GNPS platform.

## Author contributions

D.E. and E.G. conceived the study, performed the experiments, analyzed the data, and wrote the paper with inputs from the other authors. D.P. contributed bioinformatics support. C.V. performed mass spectrometry measurements. B.M. and L.M. performed NMR analyses.

## Data and materials availability

Metabolomics raw data and .mzml files were deposited on MassIVE https://doi.org/doi:10.25345/C5QB9V93Q. MSI data was deposited at metaspace https://metaspace2020.eu/project/nicotiana_msi-2022 Additional data is available on Zenodo https://doi.org/10.5281/zenodo.6536010. All scripts used in this study are available at the Github repository: https://github.com/volvox292/Nicotiana_metabolomics. All data needed to evaluate the conclusions in the paper are further present in the paper and/or the Supplementary Materials.

