## Supplementary Materials for "Evolutionary metabolomics of specialized metabolism diversification in the genus *Nicotiana* highlights allopolyploidy-mediated innovations in *N*-acylnornicotine metabolism"

**This PDF file includes:**

Supplementary Text
Figs. S1 to S15
Tables S1 to S3

**Other Supplementary Materials for this manuscript include the following:**

Data S1 to S8

#### Supplementary Text

Purification and NMR-based structural elucidation of *N*-acyl-nornicotines

*N. nesophila* leaves (354.95 g FW) were briefly rinsed with acetonitrile, the collected solvent (hereafter referred to as leaf exudate) was filtered and then concentrated under reduced pressure. The dried leaf exudate (184.4 mg) was re-dissolved in a small volume of solvent 1and loaded onto a column with 10g silica gel 60 (40-63 µm). The flash column was eluted with a gradient of petroleum ether:ethyl acetate:$NH_4OH$ from 40:60:4 (solvent 1) to 20:80:4 (solvent 2) as follows: 30 mL of solvent 1, 30 mL of solvent 1: solvent 2 (1:1) and flushing of the column with 150 mL solvent 2. 29 fractions (5-10 mL) were collected and fractions 17 to 22 and 23 to 29 were respectively combined. The latter combined fractions were further resolved by preparative HPLC with $H_2O$ (A), ACN (B) as eluents from 60 % B to 63 % B in 40 minutes on a Kinetex $C_{18}$ column (250 x 10 mm, 5 µm, 100 Å) with an injection volume of seven times of 100 µL.

Six *N*-acyl-nornicotines isolated from *Nicotiana nesophila* leaf exudates were structurally characterized by UPLC-QTOF-MS and NMR (**Fig. S11**). *N*-acyl-nornicotine **#1** (68.6 mg) was identified as 3-hydroxy-12-methyl-1-(2-(pyridin-3-yl)pyrrolidin-1-yl)tridecan-1-one and was detected as its $[M+H]^+$ adduct at *m/z* 375.3002 ($C_{23}H_{39}N_2O_2$, +1.2 ppm) in QTOF-MS. *N*-acyl-nornicotine **#2** (12.6 mg) was identified as 3-hydroxy-1-(2-(pyridin-3-yl)pyrrolidin-1-yl)tetradecan-1-one and was detected as its $[M+H]^+$ adduct at *m/z* 375.3000 ($C_{23}H_{39}N_2O_2$, +1.6 ppm). *N*-acyl-nornicotine **#3** (11.1 mg) was identified as 3-hydroxy-10-methyl-1-(2-(pyridin-3-yl)pyrrolidin-1-yl)dodecan-1-one and was detected as its $[M+H]^+$ adduct at *m/z* 361.2842 ($C_{22}H_{37}N_2O_2$, +2.0 ppm). *N*-acyl-nornicotine **#4** (9.8 mg) was identified as 3-hydroxy-12-methyl-1-(2-(pyridin-3-yl)pyrrolidin-1-yl)tetradecan-1-one and was detected as its $[M+H]^+$ adduct at *m/z* 389.3157 ($C_{24}H_{41}N_2O_2$, +1.5 ppm). *N*-acyl-nornicotines **#5** and #**6** (5.9 mg) co-eluted in the same fraction but could still be identified as 3-hydroxy-1-(2-(pyridin-3-yl)pyrrolidin-1-yl)dodecan-1-one and 3-hydroxy-10-methyl-1-(2-(pyridin-3-yl)pyrrolidin-1-yl)undecan-1-one and was detected as $[M+H]^+$ adducts at *m/z* 347.2689 ($C_{21}H_{34}N_2O_2$, +1.1 ppm) and 347.2687 ($C_{21}H_{34}N_2O_2$, +1.7 ppm). All compounds appeared as colorless oils, and seem to be present as their stereoisomers. Absolute configurations were not determined as part of this structure elucidation effort.

NMR Spectra ($^{1}H$, $^{13}C$) were performed at 298 K. 1H (500 MHz or 300 MHz) and $^{13}C$ (125 MHz) NMR chemical shifts are reported relative to residual protiated solvent. NMR are presented as follows: chemical shift (ppm), multiplicity (s = singlet, d = doublet, t = triplet, q = quartet, sept = septet, m = multiplet, br = broad), coupling constant J (Hz) and integration.

**$C_{14}$-iso-3-hydroxy-N-acyl-nornicotine (#1)**

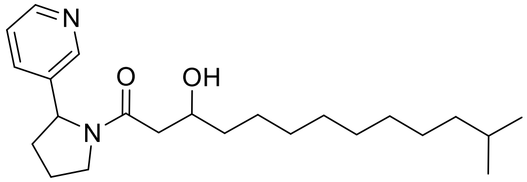

**$^{1}H$ NMR (300 MHz, $CDCl_3$) δ** = 8.61 – 8.41 (m, 2H), 7.54 – 7.43 (m, 2H), 5.20 (dd, J = 8.1, 3.1 Hz, 1H), 4.24 (s, 1H), 4.07 – 3.96 (m, 1H), 3.78 – 3.57 (m, 2H), 2.51 – 2.28 (m, 2H), 2.08 – 1.78 (m, 4H), 1.67 – 1.45 (m, 2H), 1.38 – 1.06 (m, 15H), 0.86 (d, J = 6,6 Hz, 6H).

**$^{13}C$ NMR (126 MHz, $CDCl_3$) δ** = 171.9, 148.3, 147.2, 138.4, 133.6, 123.9, 67.9, 58.7, 47.8, 41.1, 39.2, 36.6, 36.5, 33.9, 30.1, 29.8, 29.7, 28.1, 27.5, 25.7, 23.9, 22.8 (x2).

**$C_{14}$-n-3-hydroxy-N-acyl-nornicotine (#2)**

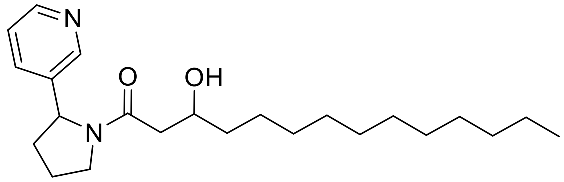

**$^{1}H$ NMR (300 MHz, $CDCl_3$) δ** = 8.59 – 8.37 (m, 2H), 7.49 – 7.41 (m, 2H), 5.20 (dd, J = 8.1, 3.1 Hz, 1H), 4.24 (s, 1H), 4.11 – 3.86 (m, 1H), 3.80 – 3.55 (m, 2H), 2.51 – 2.26 (m, 2H), 2.09 – 1.73 (m, 4H), 1.60 – 1.38 (m, 2H), 1.60 – 1.04 (m, 18H), 0.92 – 0.80 (m, 3H).

**$^{13}C$ NMR (126 MHz, $CDCl_3$) δ** = 171.9, 148.4, 147.3, 138.2, 133.5, 123.9, 68.0, 58.7, 47.8, 41.1, 36.6, 36.2, 34.0, 32.1, 29.8, 29.8, 29.7, 29.5, 25.7, 23.9, 22.8, 21.7, 14.3.

**$C_{13}$-anteiso-3-hydroxy-N-acyl-nornicotine (#3)**

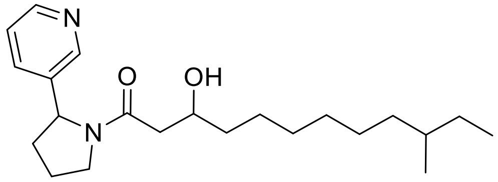

**$^{1}H$ NMR (300 MHz, $CDCl_3$) δ** = 8.58 – 8.42 (m, 2H), 7.54 – 7.40 (m, 2H), 5.20 (dd, *J* = 8.1, 3.1 Hz, 1H), 4.24 (s, 1H), 4.14 – 3.86 (m, 1H), 3.80 – 3.56 (m, 2H), 2.51 – 2.25 (m, 2H), 2.10 – 1.74 (m, 4H), 1.59 – 1.42 (m, 2H) 1.40 – 0.98 (m, 13H), 0.92 – 0.76 (m, 6H).

**$^{13}$C NMR (126 MHz, $CDCl_3$) δ** = 171.9, 148.4, 147.3, 138.2, 133.5, 123.9, 67.9, 58.7, 47.7, 41.1, 36.7, 36.6, 34.5, 33.9, 30.1, 29.8, 29.6, 25.7, 23.9, 21.7, 19.4, 11.6.

**$C_{15}$-anteiso-3-hydroxy-N-acyl-nornicotine (#4)**

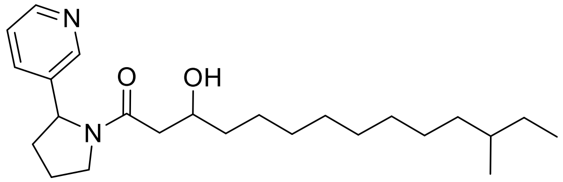


**$^1$H NMR (300 MHz, CDCl3) δ** = 8.74 – 8.33 (m, 2H), 7.51 – 7.41 (m, 2H), 5.20 (dd, J = 8.1, 2.9 Hz, 1H), 4,24 (s, 1H), 4.12 – 3.86 (m, 1H), 3.82 – 3.54 (m, 2H), 2.51 – 2.27 (m, 2H), 2.08 – 1.70 (m, 4H), 1.60 – 1.44 (m, 2H), 1.42 – 0.97 (m, 17H), 0.91 – 0.77 (m, 6H).

**$^{13}$C NMR (126 MHz, $CDCl_3$) δ** = 171.9, 149.6, 147.5, 138.8, 132.9, 124.1, 68.0, 58.8, 47.8, 41.1, 36.8, 36.6, 36.3, 34.5, 34.0, 30.1, 29.8, 29.6, 27.2, 25.7, 23.9, 21.7, 19.4, 11.6.

**$C_{12}$-n-3-hydroxy-N-acyl-nornicotine (#5)**

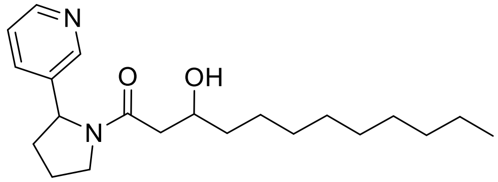


**$^1$H NMR (300 MHz, CDCl3) δ** = 8.61 – 8.39 (m, 2H), 7.51 – 7.42 (m, 2H), 5.20 (dd, J = 8.1, 3.1 Hz, 1H), 4.24 (s, 1H), 4.10 – 3.87 (m, 1H), 3.81 – 3.57 (m, 2H), 2.51 – 2.28 (m, 2H), 2.07 – 1.67 (m, 4H), 1.61 – 1.44 (m, 2H), 1.42 – 1.04 (m, 14H), 0.90 – 0.84 (m, 3H).

**$^{13}$C NMR (126 MHz, $CDCl_3$) δ** = 172.6, 148.3, 147.3, 138.2, 132.9, 123.6, 68.0, 58.7, 47.8, 41.1, 39.2, 36.3, 32.0, 29.8, 29.5, 27.5, 25.7, 23.9, 22.8, 21.7, 14.3.

**$C_{12}$-iso-3-hydroxy-N-acyl-nornicotine (#6)**

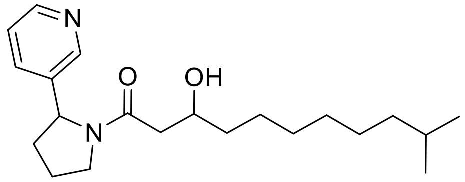


**$^1$H NMR (300 MHz, CDCl3) δ** = 8.61 – 8.39 (m, 2H), 7.51 – 7.42 (m, 2H), 5.20 (dd, J = 8.1, 3.1 Hz, 1H), 4.24 (s, 1H), 4.10 – 3.87 (m, 1H), 3.81 – 3.57 (m, 2H), 2.51 – 2.28 (m, 2H), 2.07 – 1.67 (m, 4H), 1.61 – 1.44 (m, 2H), 1.42 – 1.04 (m, 11H), 0.86 (d, J = 6.6 Hz, 6H).

**$^{13}$C NMR (126 MHz, $CDCl_3$) δ** = 171.9, 149.2, 147.6, 138.4, 133.6, 123.9, 68.0, 58.7, 47.8, 41.1, 39.2, 36.6, 33.9, 30.0, 29.7, 28.1, 27.5, 25.7, 23.9, 22.8 (x2).

Creating an *in silico* MS/MS library for approximately 1 million natural products and optimizing its rapid interrogation

To increase the fragmentation-based annotation rate of our MS/MS dataset and cover compound classes for which experimental high-resolution MS/MS spectra are scarcely present in databases, we created an *in silico* spectral library for about 1.1 million chemical structures corresponding to natural products. Such an approach has initially been pioneered by (*42*), but with chemical entries (~ 220 000) retrieved from the copyrighted Dictionary of Natural Products. Here, structures were downloaded from several public chemical libraries (Table **S3**) and converted to InChI format (Script **S1**, see Supplementary Text "**Code Availability and description"**). Individual databases were parsed and duplicates were removed (Scripts **S2** & **S3**, see Supplementary Text "**Code Availability and description"**), which resulted in 1,103,179 structures. In order to be submitted in a highly parallelized manner at the High Performance Computing Center of the University of Strasbourg, the concatenated structure database was split randomly into 1103 parts for *in silico* fragmentation using CFM-predict (version 4.0.8) (23). *In silico* MS/MS spectra generated for different collision energies (10 eV, 20 eV and 40 eV) were merged to create a final set of 1,066,512 composite spectra (Script **S5**, see Supplementary Text "**Code Availability and description"**). Interrogation of this database was implemented with an "optimized version" of the MatchMS pipeline using the Spec2vec score (*75*) (Scripts **S6** & **S7**, see Supplementary Text "**Code Availability and description"**) or the modified cosine score (S**26**, S**27**). The Spec2vec search took 20 hours with 150 Gb RAM on 1 core with a retrained model that included the spectra of our *in silico* database. The modified cosine score search was more computationally intensive and took 3.5 days parallelized on 22 cores and 350 Gb of RAM. The database is available through the GNPS environment and Zenodo link: https://doi.org/10.5281/zenodo.6536010

Comparison of hits retrieved from *in silico* spectral interrogations using the Jassbi *Nicotiana* chemical database

A small-scale attempt at comparing the performance of the *in silico* fragmentation tools was conducted using the Jassbi *Nicotiana* chemical database (*32*) (**Fig. S7**). The numbers of annotations were 999 with CFM-ID v4.0 and the modified cosine (score above 0.5 and more than 5 matching peaks), 65 significant annotations with Moldiscovery, 159 annotations with QCxMS and the modified cosine (score above 0.5 and more than 5 matching peaks). Numbers of annotations were compared and also their classifications retrieved with NP-classifier, all the tools show terpenoids were the most common compound class (**Fig. S6**). The singularity image of the simplified batchmode used for running QCxMS is available at the following Zenodo link: https://doi.org/10.5281/zenodo.6536010 .

Code availability and description

All scripts used in this study are available at the Github repository: https://github.com/volvox292/Nicotiana_metabolomics

**S1** openbabel_conversion.ipynb | *Converts Smiles to InchI for the creation of the* in silico *spectral database*

**S2** reformater.R | *Parsing/cleaning of the structure databases for the creation of the* in silico *spectral database*

**S3** add_openbabel_info.R | *Structure database merging and duplicate removal for the creation of in silico spectral database*

**S4** run_cfmid_mesocenter.sh & cfmid_commands.txt | *Runs CFM-ID on the University Strasbourg HPC Cluster*

**S5** Process_mgf.ipynb | *Creates composite spectra from merging spectra obtained by CFM-ID at different collision energies*

**S6** matchms_spec2vec.py | *Interrogation of the* in silico *spectral database using spec2vec as scoring metric*

**S7** matchms_scores_analysis.py | *Used for Database Matching of CFM ID on HPC Cluster*

**S8** MatchMS-v1-cosine-msp.ipynb | *Used for Database Matching of Nicotiana DB*

**S9** MatchMS-v1-cosine.ipynb | *Used for Database Matching of Jassbi*

**S10** Batch-QTOF-sens-v3.xml | *Used for Batch Mode processing of the Dataset*

**S11** mgf-rem-redundancy-v4.ipynb | *Remove redundant features*

**S12** Sirius-removev2.ipynb | *Remove redundant IDs from Sirius mgf file*

**S13** run_sirius.sh | *Used to run Sirius on HPC Cluster*

**S14** degree-unsaturation-sirius.ipynb | *Restore Feature ID from Sirius ID and Calculate degree of unsaturation, requires molmass package*

**S15** compound-id-sirius.ipynb | *Restore Feature ID from Sirius ID*

**S16** canopus_consensus_ms2lda.ipynb | *Merge the outputs of all the tools into one big table also get consensus substructures (based on ms2lda motifs) for insilico-tools and propagate canopus within networks*

**S17** MSLDAmerge-motfs.ipynb | *Get Motifcount based on Presence of Feature*

**S18** MSLDAmerge-motfs-sumall.ipynb | *Get Motifcount based on Presence of Feature within all Tissues*

**S19** canopus_consensus.ipynb | *Script to merge the outputs of all the tools into one big table also get consensus substructures for insilico-tools and propagate canopus within networks*

**S20** phylometabo.ipynb | *Calculate Pairwise Distance Matrix based on Data of Motif Figure and Network Figure*

**S21** phylo.Rmd | *Plot Phylogenies from pairwise Distance Matrix based on APE package*

**S22** sum_molformula_areas.ipynb | *Sum Areas of NANNs based on identical Molecular Formula*

**S23** nann_bubbles.ipynb | *Sum Areas based on Carbon Chain of NANNs, split by hydroxylation or not*

**S24** Networkclustermap.ipynb | *Sum all areas of Networks per Samples*

**S25** ASR-single.Rmd | *Ancestral State reconstruction based on MBASR*

**S26** dbsearch.py | *Script to run modified cosine score based search on big in-silico db*

**S27** run_db.py | *Script to run modified cosine score based search on big in-silico db*

**S28** group_for_treemap.ipynb | *Used to group and sum canopus classes peak areas*

**S29** alpha_diversity.ipynb | *Calculate alpha diversity based on shannon entropy*

**S30** Vegan_calculations.Rmd | *NMDS using vegan package*

**S31** cosine_distance_sp_canopus.ipynb | *Calculate distances between species and canopus classes*

**Fig. S1.**

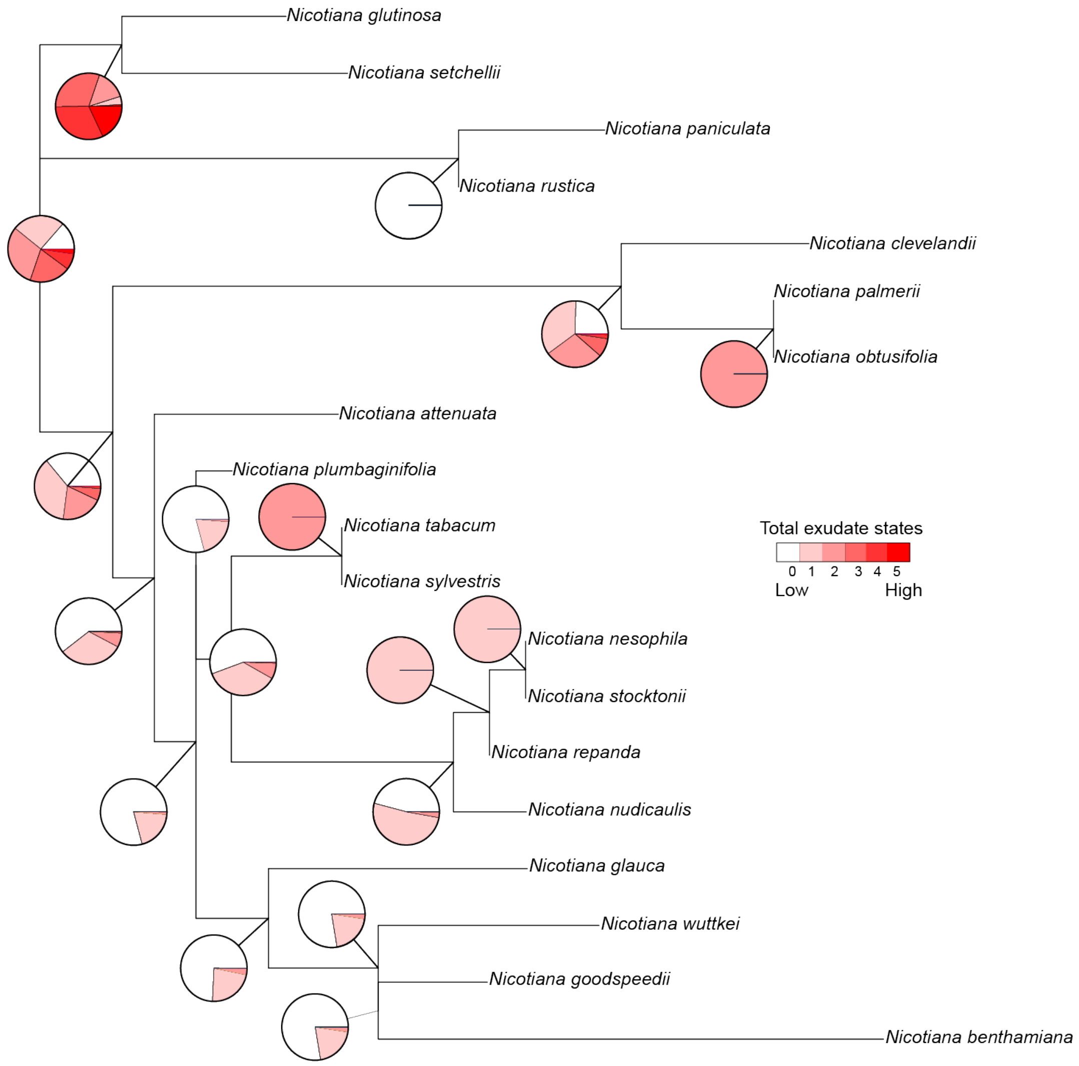


**Fig. S1. Ancestral trait reconstruction on the absolute amount of exudates collected from the focal species.** Total leaf exudates' dry weights (**Table S1**) from the focal species was transposed as relative scaling into an ordered trait (total exudate states colored from white to dark red) and used as input for ancestral state reconstruction using the MBASR software with default settings. The species tree was constructed from *matK* gene sequence as described in (*71*).

**Fig. S2.**

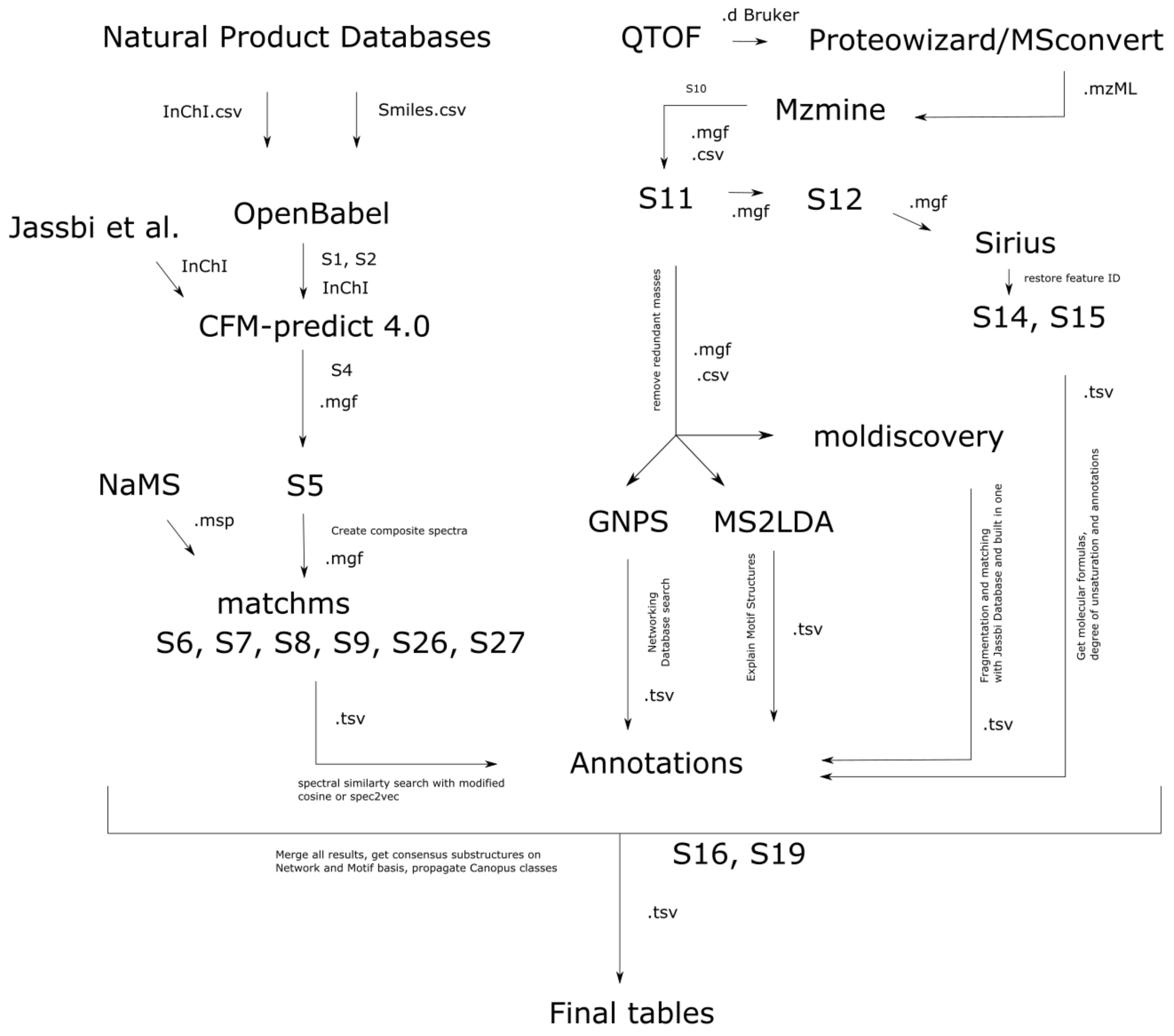


**Fig. S2. Architecture of the metabolomics data processing workflow with reference to custom scripts developed for this study.** All referred scripts (See Supplementary Text "**Code description and availability**") are available at the Github repository: https://github.com/volvox292/Nicotiana_metabolomics

**Fig. S3.**

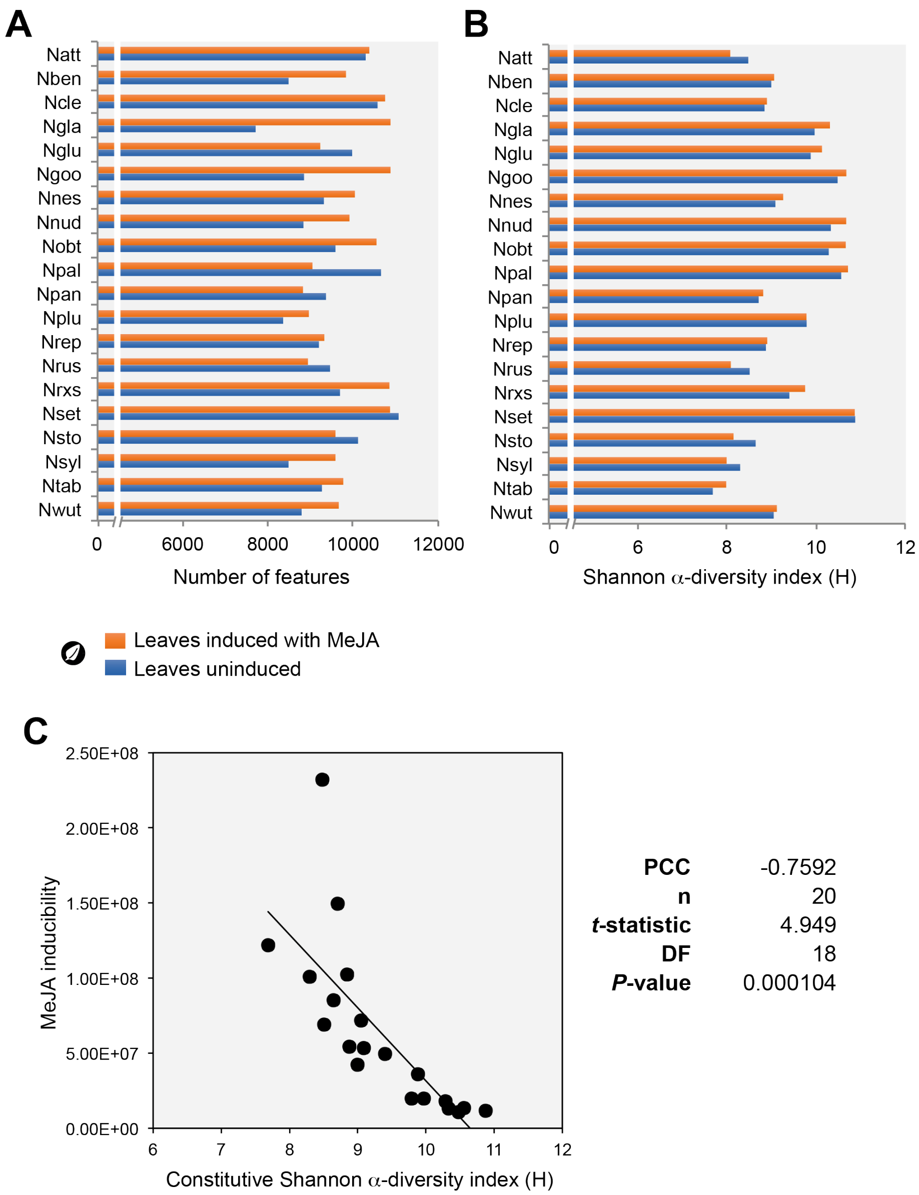


**Fig. S3. Comparison of diversity analysis of profiles of uninduced and MeJA-induced leaves. (A)** Bar chart depicting numbers of features detected *per* species metabolic profiles. **(B)** bar chart depicts the Information Theory Shannon α-diversity (H) as an index of feature richness. *Nicotiana* species (see **Table S1** for complete species information) are alphabetically-ordered. **(C)** Biplot visualizing the inter-species negative correlation between leaf metabolome MeJA inducibility (calculated from the Euclidean distance between MeJA-induced and uninduced leaf profiles) and constitutive (uninduced leaf samples) α-diversity. PCC, Pearson Correlation Coefficient; DF, Degree of Freedom.

**Fig. S4.**

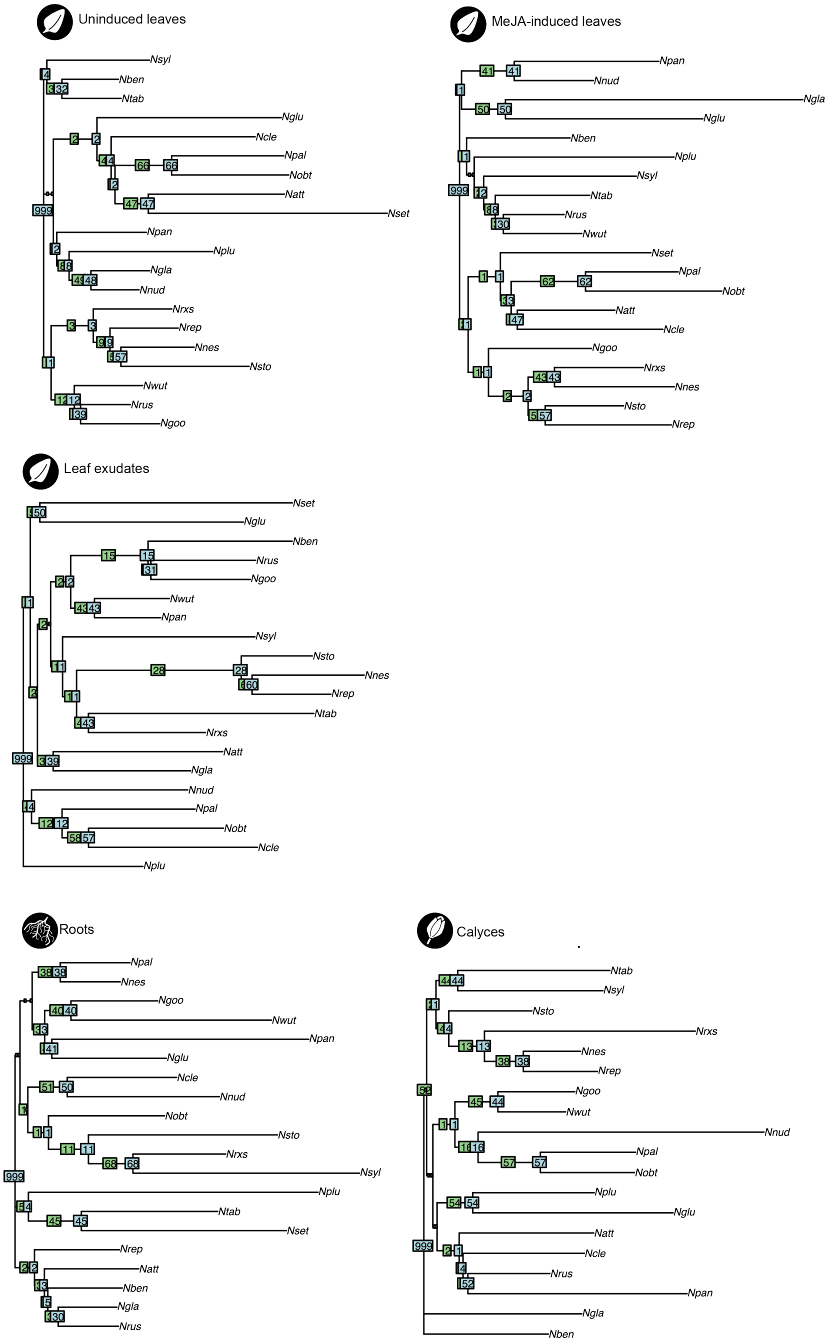


**Fig. S4. Tissue-type "phylometabolomics" tree computed from the molecular networking information.** To analyze the relatedness of species' metabolomes, we first computed inter-species Euclidean distances based on the molecular networking information and used the resulting distance matrices for constructing "phylometabolomics" trees based on the Neighbor-Joining algorithm (bootstrap values derived from 999 iterations).

**Fig. S5.**

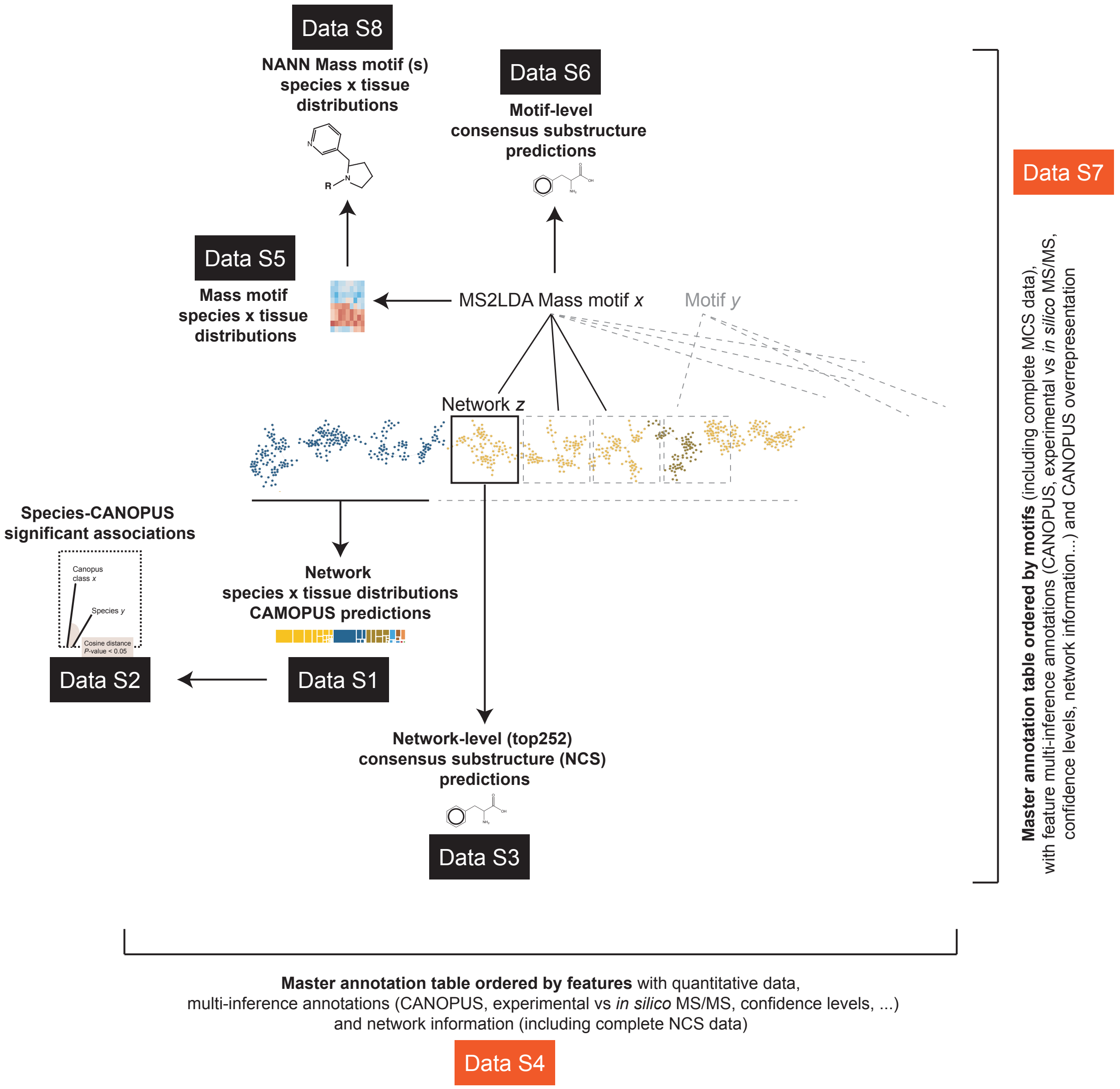


**Fig. S5. Overview of Supplementary Data**

**Fig. S6.**

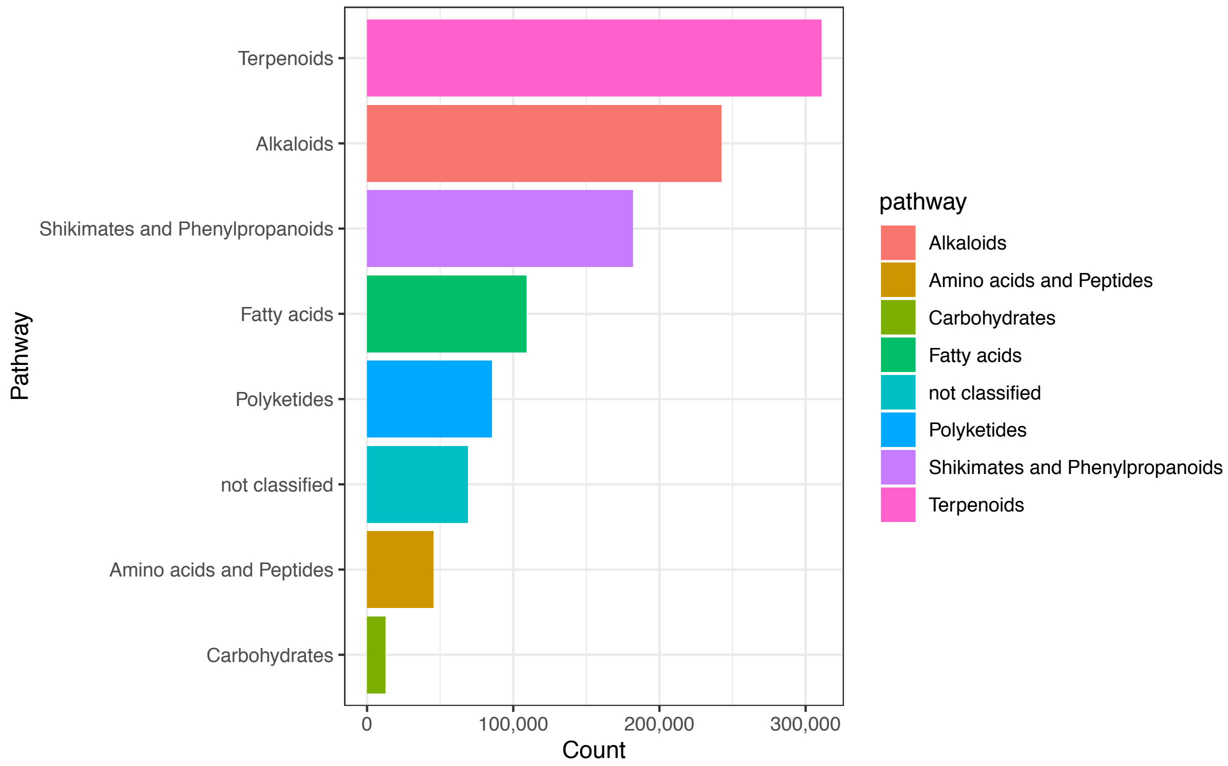


**Fig. S6. Composition of the 1 million *in silico* natural products library.** Chemical entries were classified by NP-classifier. The creation of this in silico spectral database is referred in the Supplementary Text **"Creating an *in silico* MS/MS library for approximately 1 million natural products and optimizing its rapid interrogation".** The list of natural product database employed is presented in **Table S2**.

**Fig. S7**

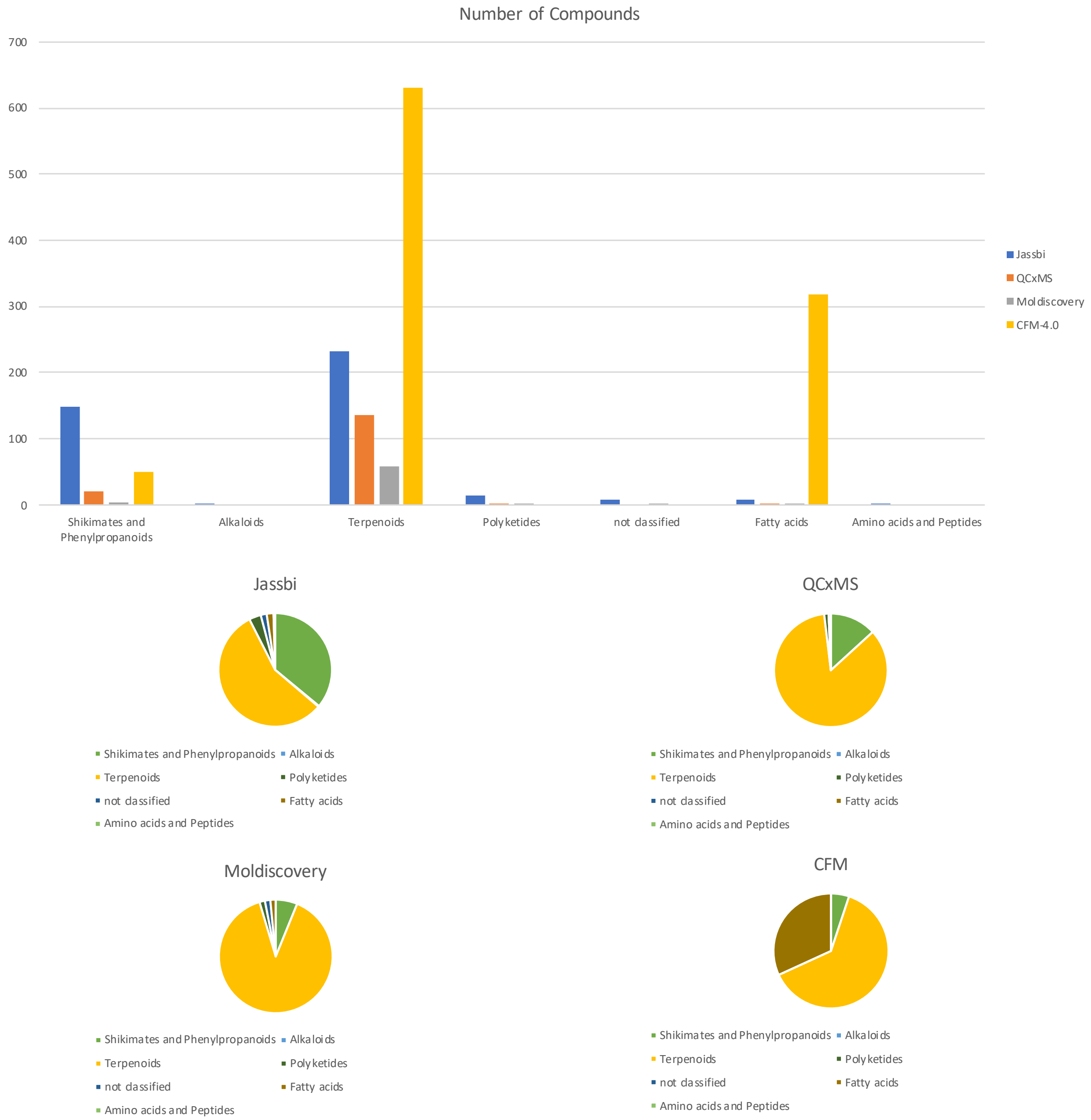


**Fig. S7. Comparison, using NP-classifier, of hit chemical classes retrieved from *in silico* spectral interrogations using the Jassbi *Nicotiana* chemical database.** Top panel, number of compound hits *per* retrieved from searching the *Nicotiana* dataset against the Jassbi *Nicotiana* chemical database (*32*) with the different tools. The numbers of annotations were 999 with CFM-ID v4.0 and the modified cosine (score above 0.5 and more than 5 matching peaks), 65 significant annotations with Moldiscovery, 159 annotations with QCxMS and the modified cosine (score above 0.5 and more than 5 matching peaks). Lower panel, hits were then classified with NP-classifier (pie charts in the lower panel).

**Fig. S8.**

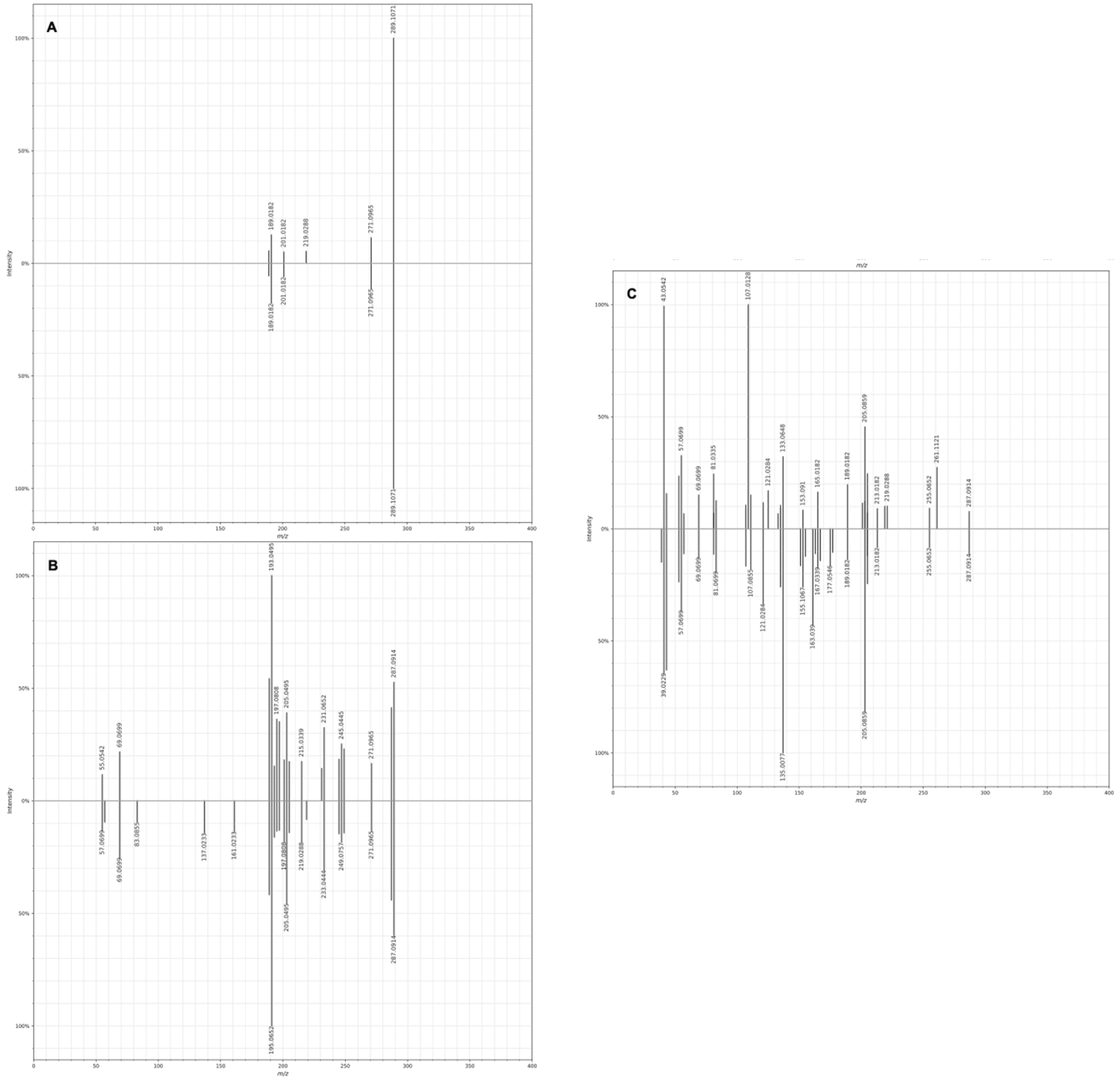


**Fig. S8. Predictions for (+)- and (-)-shikonin MS/MS spectra.** Mirror plots of (+)-shikonin (top) and (-)-shikonin (bottom) *in-silico* fragmentation spectra created with CFM 4.0. **(A)** low collision energy (10eV), **(B)** medium collision energy (20 eV), **(C)** high collision energy (40 eV). Slight variations in peak intensity but also appearance of additional peaks can be observed when comparing the two stereoisomers.

Fig. S9

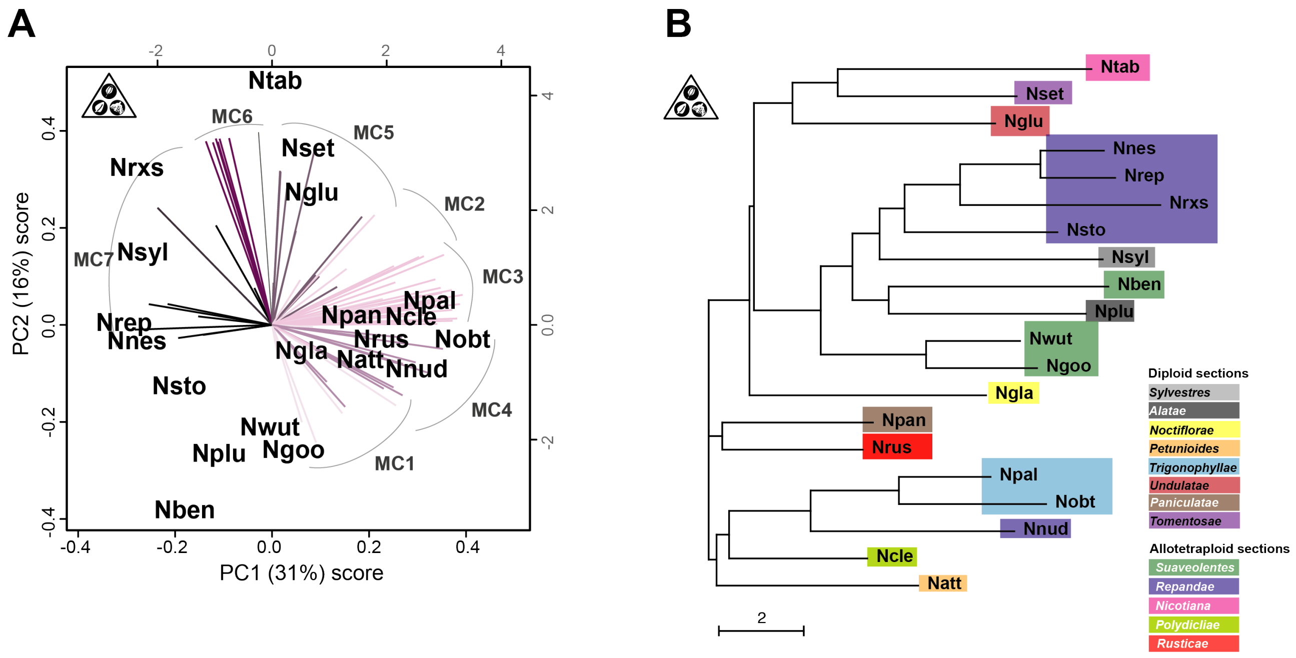


**Fig. S9. The top76 MS2LDA mass motifs capture the structural diversity among *Nicotiana* phylogenetics section. (A)** Score plot obtained from the principal component analysis (2 first PCs) of species-level MS motif counts. Loadings exerted on sample PC coordinates are mapped for each MS motifs. Colors refer to motif clusters (MC) derived from the hierarchical clustering analysis based on the species-level motif-associated peak intensity data (Z-score normalized) form top76 mass motifs inferred by unsupervised decomposition of overall MS spectra via the text-mining program MS2LDA from the whole-tissue data (**Figure 5**). **(B)** Phylogenetic tree (pairwise euclidian distances plotted as neighbor joining tree) top76 mass motif data indicating the power of mass motif analysis as a data reductionality approach to compare whole-tissue species-level metabolomes. *Nicotiana* phylogenetic sections are color-coded.

### Fig. S10

## A

| | CANOPUS Super-class | CANOPUS Sub-class | CANOPUS Most-specific class | Network-level consensus substructure (NCS) NCS 0 | NCS 1 | NCS 2 | NCS 3 |
|---|---|---|---|---|---|---|---|
| **Network #1043** Kaurane-based diterpenes enriched in Ntab, Nglu, Nset, Nsyl leaf exudates | Lipids and lipid-like molecules | Diterpenoids | Diterpenoids | CC1CCCC2(C)C1CCC13CCC(CCC12)C3 | C=C1CCC2C(C)(CO)CCCC2(C)C1 | CC=CC=CCCC | |
| **Network #262** Diterpenes enriched in Nsyl, Ntab, Nrxs, Nset leaf surfaces and calyces | Lipids and lipid-like molecules | Diterpenoids | Diterpenoids | CC=C(C)C | CCCCC=C(C)C | CC=C(C)C | |
| **Network #468** Hydroxycinnamoyl-spermidines enriched in Ngla calyces and MeJA-induced leaves | Organic acids and derivatives | Amino acids, peptides, and analogues | Amino acids and derivatives | NC(=O)C=CC1:C:C:C(O):C(O):C:1 | C=CC1:C:C:C(O):C(O):C:1 | NC(CC1:C:C:C:C:C:1)C(=O)O | OC1:C:C:C:C:C:1O |
| **Network #1662** Saccharolipids (O-acyl sugars) enriched in Npal, Nobt, Ncle, Nglu, Nnud leaf exudates/calyces | Lipids and lipid-like molecules | Fatty acyl glycosides | Saccharolipids | CCCCCC(=O)OCCC(O)C(O)CO | COC1OC(CO)C(O)C(O)C1O | CC=CC=CCCCCCC | |
| **Network #222** Saccharolipids (O-acyl sugars) enriched in Nben, Nrus, Ngoo, Nwut leaf exudates/calyces | Lipids and lipid-like molecules | Fatty acyl glycosides | Saccharolipids | OCC1OC(OC2(CO)OCC(O)C2O)C(O)C(O)C1O | CC=CCCCCCC(=O)OC | C=CCCCCC | |
| Figure 5: Motif Strepsalini_110 Network #721 | Lipids and lipid-like molecules | Diradylglycerols | 1,2-diacylglycerols | CCCCCCCC | CCCC | CC(=O)OC(C)CO | CCCCCCCCCCCCCC=O |
| Figure 5: Motif GNPS_37 Network #486 | Organic oxygen compounds | Carbohydrates and carbohydrate conjugates | Phenolic glycosides | CC=CC1:C:C:C(O):C(O):C:1 | COC1OC(CO)C(O)C(O)C1O | CC1:C:C:C:C:C:1 | |

## B

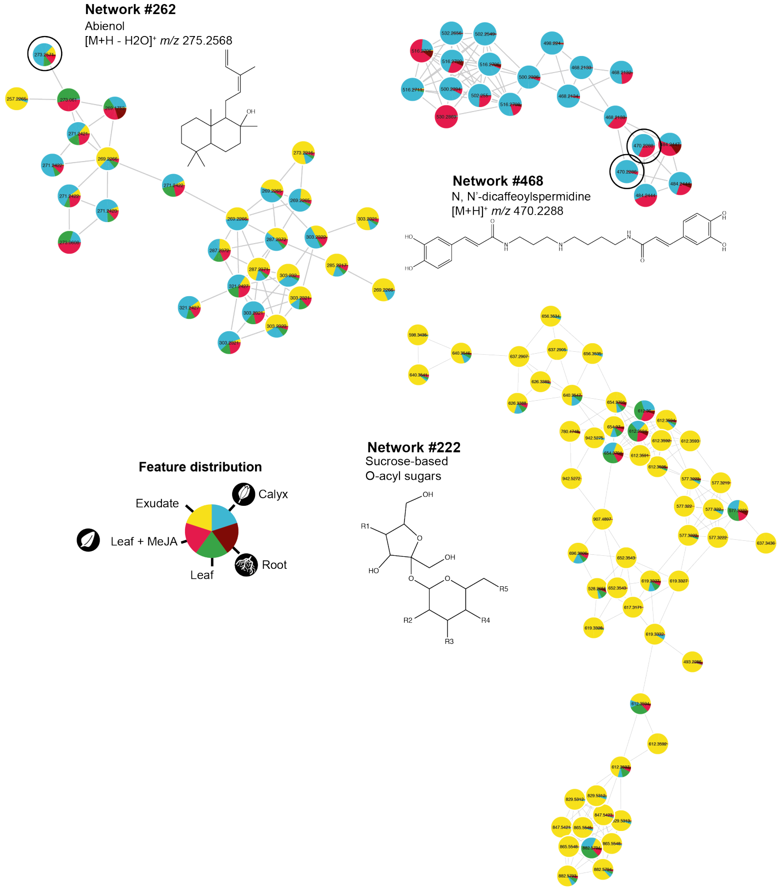

**Fig. S10. Network consensus substructure analysis for selected networks. (A)** Network consensus substructure analysis for selected network for which in silico MS/MS annotations were collected (see **Data S1** for NCS of the top252 networks) and which corresponded to CANOPUS predictions specifically associated with certain species (**Data S2**). Top4 NCS are provided as well as comments on manual annotation, species association. NCS for two networks exemplified in **Figure 5** are further presented. **(B)** Molecular networks for networks #262, #468 and #222, corresponding respectively to labdane diterpenes, hydroxycinnamoyl-spermidines and sucrose-based *O*-acyl sugars. Representative high confidence annotation metabolites are presented. Note that *O*-acyl sugars, only the sucrose core structure is presented. R1-to-R5 moities refer to H or branch or straight short-to-medium fatty acyl chains as described in (*76*). Node colors denote for the species-overall feature relative abundance in the analyzed tissues. Complete data are accessible in **Data S3** and **S4**.

**Fig. S11**

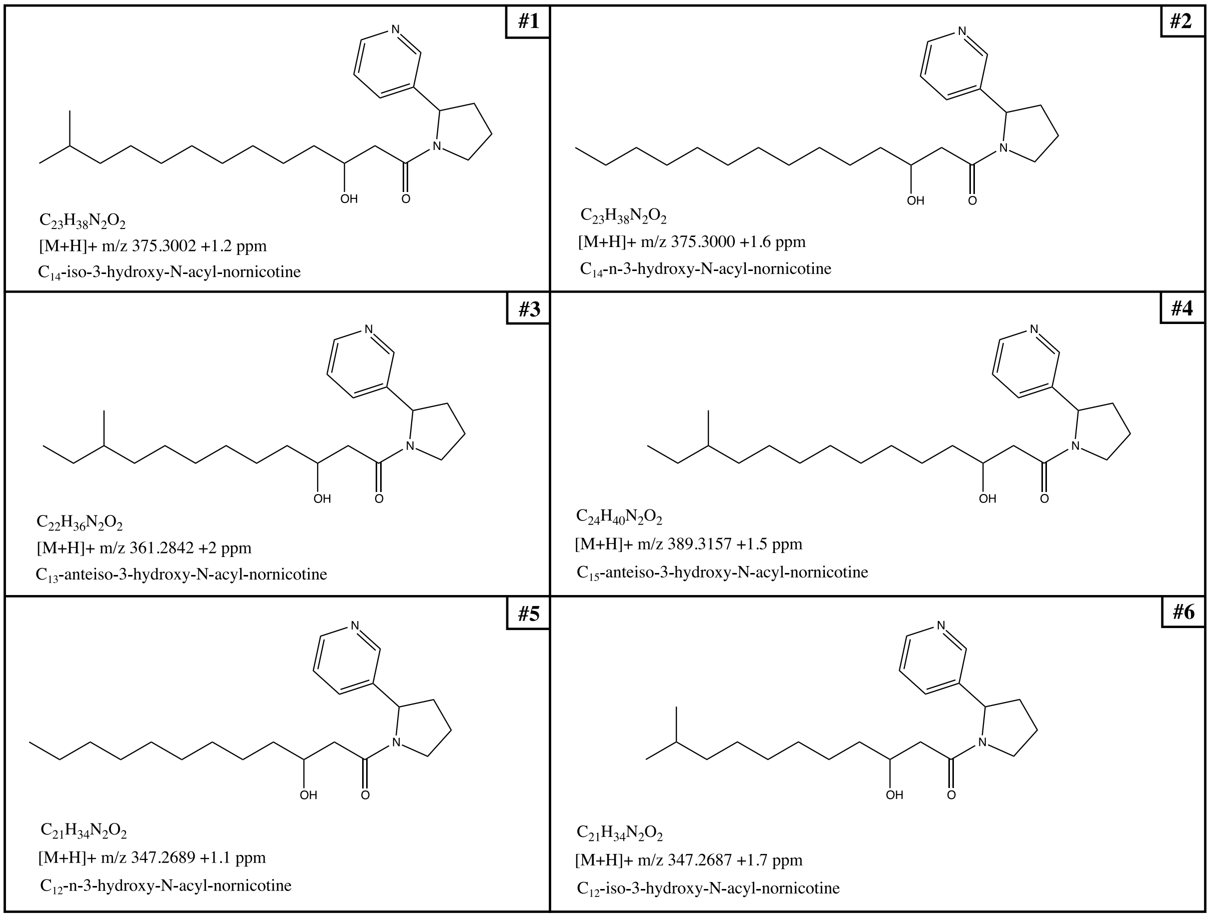


**Fig. S11. NMR structure elucidated as NANNs from leaf surface exudates of *N. nesophila*.** The purification by column chromatography and preparative HPLC yielded: compound #1 68.6 mg, #2 12.6 mg, #3 11.1 mg, #4 9.8 mg, compound #5 and #6 5.9 mg co-eluted in the same fraction but could still be elucidated (See Supplementary Text "**Purification and NMR-based structural elucidation of *N*-acyl-nornicotines**").

Fig. S12

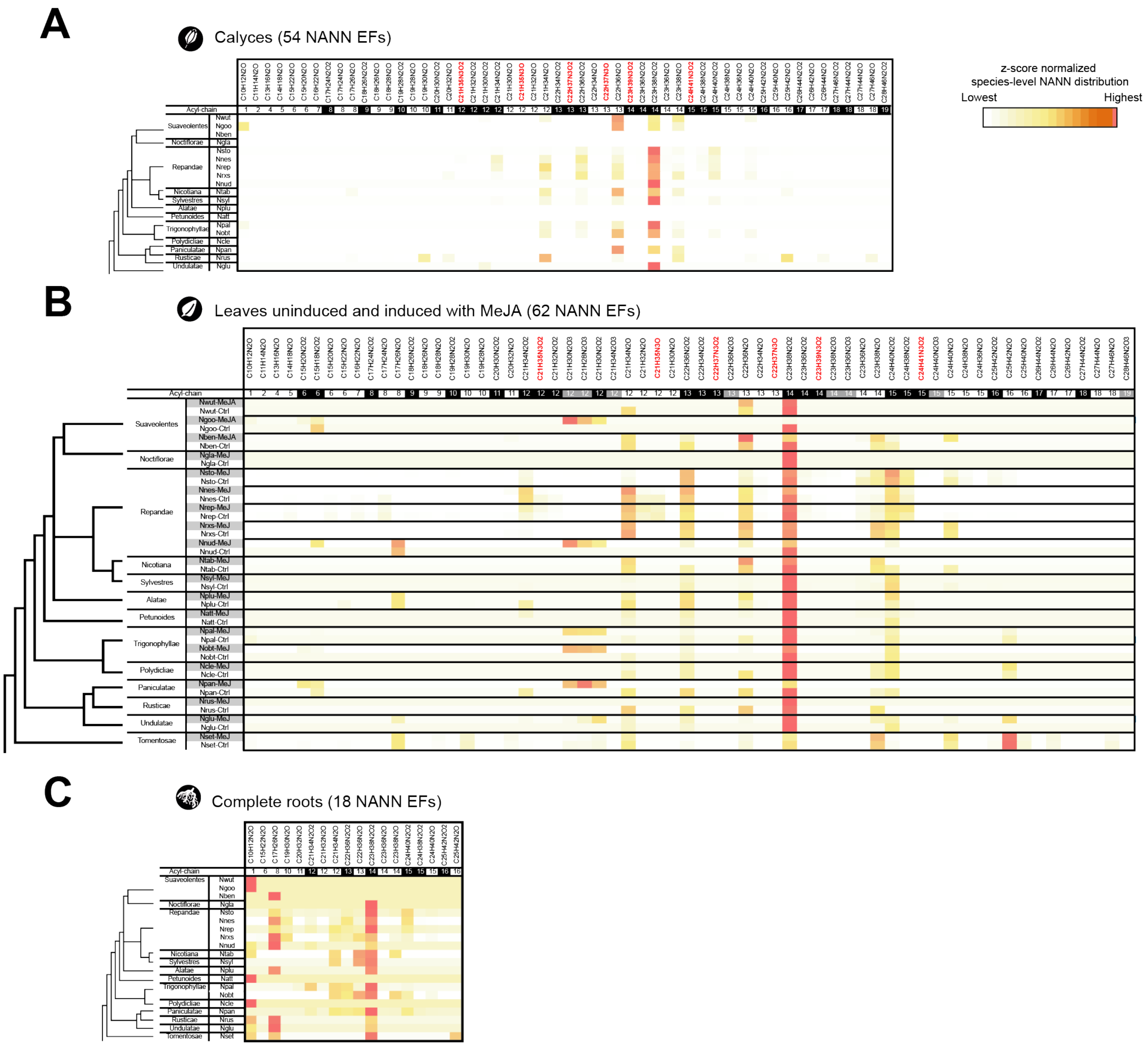


**Fig. S12. Species-level NANN elemental formula distribution in calyces (A), uninduced/MeJA-induced leaves (B) and in roots (C).** Heatmaps depicts Z-score normalized species-level NANN distributions. Acyl chain information data provides indication on the acyl chain length and of its 3-hydroxylation. Elemental formulas indicative of non-canonical data are highlighted: in red, for $N_3$ NANNs, black and grey cells in the acyl chain lines refer to mono-hydroxylated and di-hydroxylated fatty acyl chain respectively. Grey cells in the species column refer to MeJA treatment for panel (B). Complete NANN data are accessible in **Data S8**.

**Fig. S13**

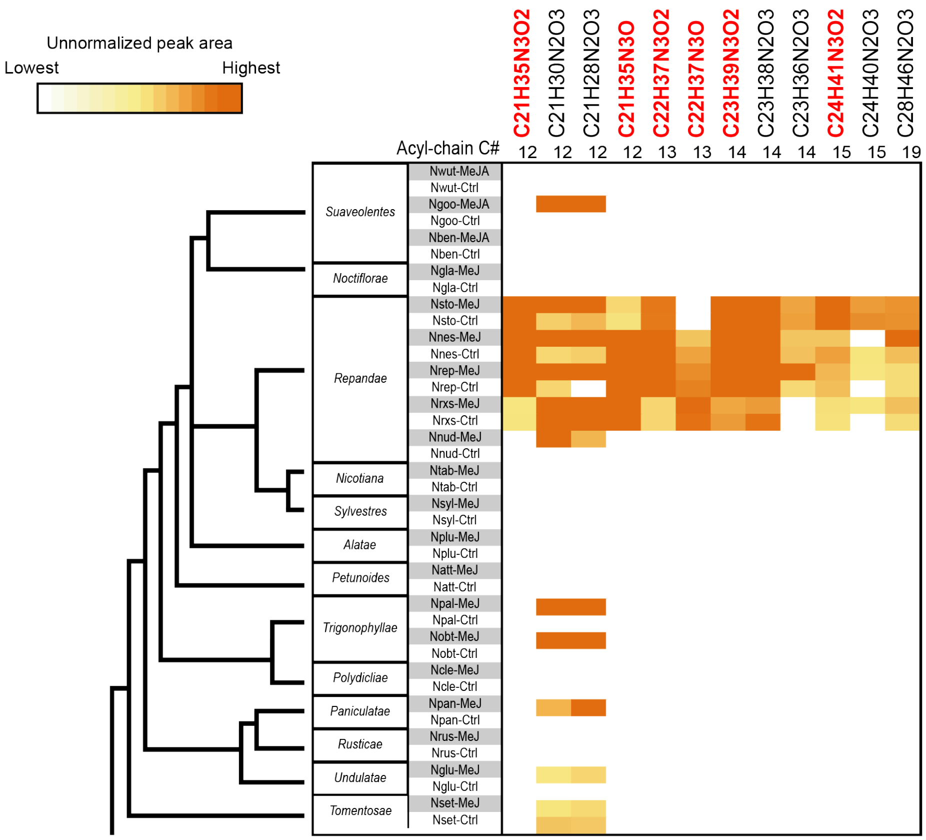


**Fig. S13. Non-canonical NANNs are dominantly detected in complete leaves of the *Repandae*.** Non-canonical NANN elemental formulas harbor three atoms of O or N (in red), respectively indicative of a second hydroxyl function or of an amine within the acyl chain (**Figure 7**). 3N-containing NANNs mostly specific from leaf tissues (**Data S8**). Heatmap depicts unnormalized peak areas for $[M+H]^+$ adducts of each NANN.

**Fig. S14**

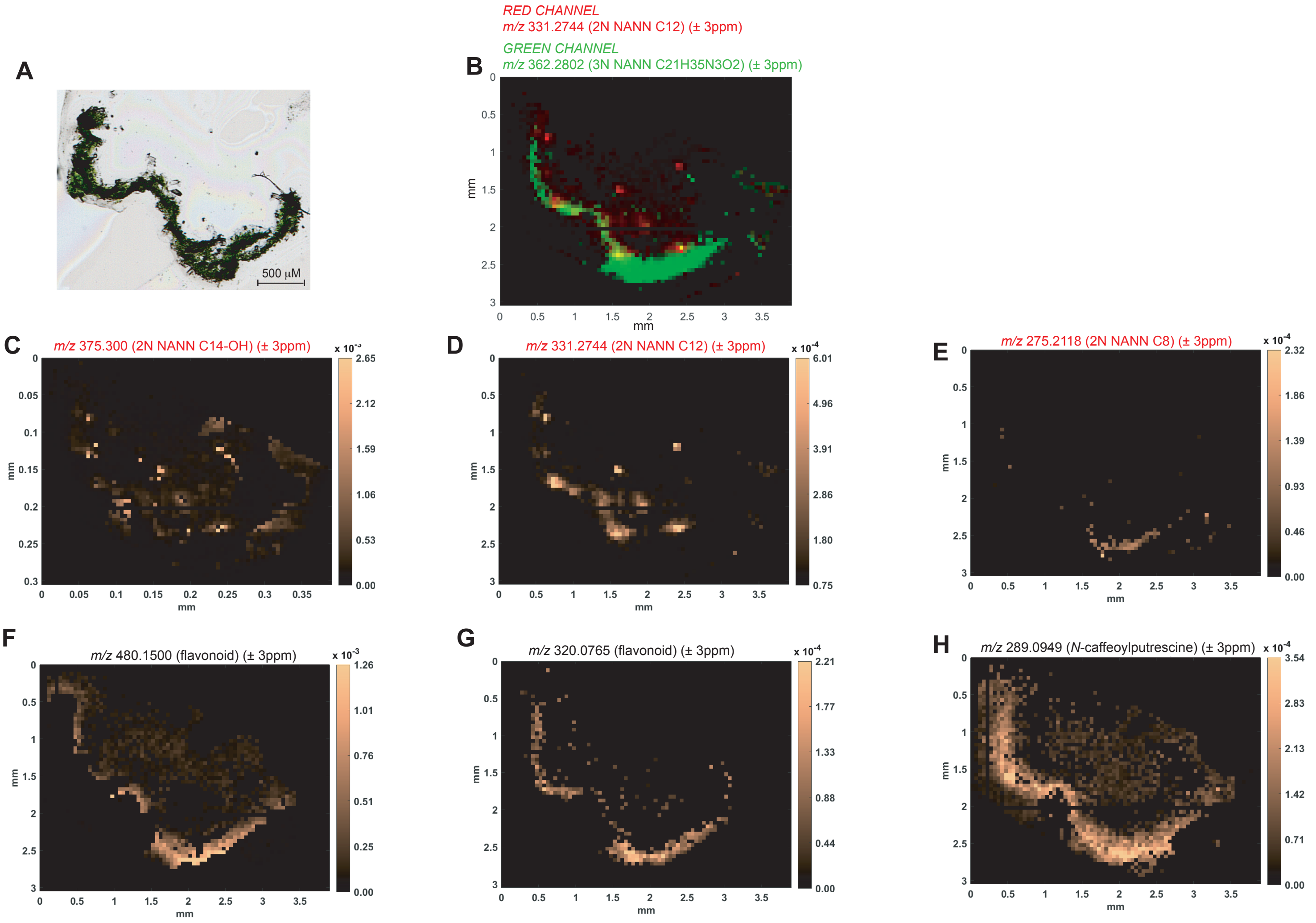


**Fig. S14. MALDI MS images depicting spatially-resolved relative abundance of selected metabolites in a leaf cross section of *Nicotiana nesophila.*** **(A)** Optical image of the matrix-embedded leaf cut used for MALDI MSI. **(B)** Overlay of MSI data for m/z 331.2744 (± 3ppm) (Red channel, "canonical" $N_2$ NANN with a $C_{12}$ acyl chain) exhibiting spot-like distribution reflecting its trichome exudation as droplets and for m/z 362.2802 (Green channel, "non-canonical" $N_3$ NANN) exhibiting a quasi-uniform distribution within the complete leaf section (mostly matching that of well-characterized leaf flavonoids, **F-I**). **(C-E)** Selected *m/z* signals for three "canonical" $N_2$ NANN also exhibiting spot-like distribution presence as spots reflecting their trichome exudation as droplets. **(F-H)** Selected *m/z* signals for two predicted flavonoids and *N*-caffeoylputrescine.

#### Fig. S15

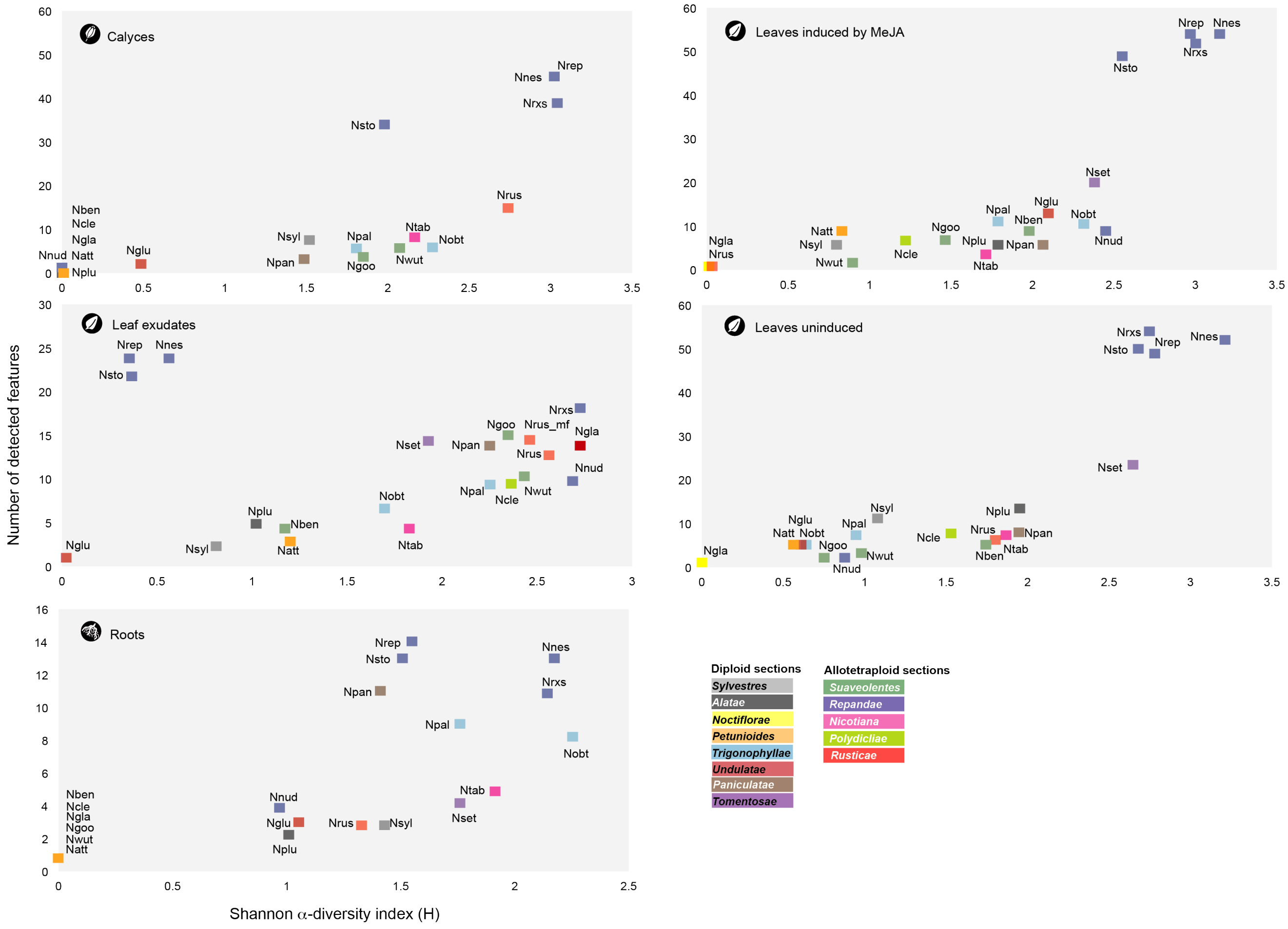


**Fig. S15. Diversity analysis of tissue-level NANN profiles.** Biplots depict the number of NANN features and the Information Theory Shannon α-diversity as an index of NANN richness *per* tissue. *Nicotiana* phylogenetics sections are color-coded.

**Table S1**

| Abbreviation | Species section | Source | Section | Ploidy | Maternal Progenitor | Paternal Progenitor | Exudate (mg) | Exudate (mg/g) |
|---|---|---|---|---|---|---|---|---|
| Ntab | *Nicotiana tabacum* | IBMP seed collection | *Nicotiana* | Allotetraploid | *N. sylvestris* | *N. tomentosiformis* | 37.7 | 1.0 |
| Ncle | *Nicotiana clevelandii* | IBMP seed collection | *Polydicliae* | Allotetraploid | *N. obtusifolia* | *N. attenuata* | 18.7 | 0.6 |
| Nnes | *Nicotiana nesophila* | Nijmegen, 96475009 | *Repandae* | Allotetraploid | *N. sylvestris* | *N. obtusifolia* | 2.2 | 0.3 |
| Nnud | *Nicotiana nudicaulis* | USDA, PI 555540 | *Repandae* | Allotetraploid | *N. sylvestris* | *N. obtusifolia* | 7.8 | 0.2 |
| Nrep | *Nicotiana repanda* | IPK, NIC5 | *Repandae* | Allotetraploid | *N. sylvestris* | *N. obtusifolia* | 148.6 | 0.6 |
| Nrxs | *Nicotiana repanda x sylvestris* | Bergerac, 550 | *Repandae* | Allotetraploid | Hybrid *N. repanda* x *N. sylvestris* | | 17.7 | 0.4 |
| Nsto | *Nicotiana stocktonii* | IPK, NIC29 | *Repandae* | Allotetraploid | *N. sylvestris* | *N. obtusifolia* | 541.4 | 0.6 |
| Nrus (Nrusmf) | *Nicotiana rustica* | IBMP seed collection | *Rusticae* | Allotetraploid | *N. paniculata* | *N.undulata* | 0.5 (Nrusmf, 215.6) | 0.02 (Nrusmf, 2.7) |
| Nben | *Nicotiana benthamiana* | IBMP seed collection | *Suaveolentes* | Allotetraploid | | | 12.9 | 0.6 |
| Ngoo | *Nicotiana goodspeedii* | Bergerac, 1129 | *Suaveolentes* | Allotetraploid | *N. sylvestris, sections Noctiflorae and Petunioides* | | 0.9 | 0.04 |
| Nwut | *Nicotiana wuttkei* | Bergerac, 1114 | *Suaveolentes* | Allotetraploid | | | 1.2 | 0.03 |
| Ngla | *Nicotiana glauca* | IBMP seed collection | *Noctiflorae* | Diploid (homoploid hybrid) | *Sections Noctiflorae and Petunioides* | | 1.3 | 0.04 |
| Nglu | *Nicotiana glutinosa* | Bergerac, 631 | *Undulatae* | Diploid (homoploid hybrid) | *Sections Tomentosae and Undulatae* | | 64.5 | 2.5 |
| Nplu | *Nicotiana plumbaginifolia* | IBMP seed collection | *Alatae* | Diploid | | | 5.9 | 0.2 |
| Npan | *Nicotiana paniculata* | Bergerac, 522 | *Paniculatae* | Diploid | | | 1.0 | 0.02 |
| Natt | *Nicotiana attenuata* | ITB (MPI-ICE), Utah acc. | *Petunioides* | Diploid | | | 7.3 | 0.3 |
| Nsyl | *Nicotiana sylvestris* | IPK, NIC37 | *Sylvestres* | Diploid | | | 72.1 | 1.3 |
| Nset | *Nicotiana setchellii* | Bergerac, 644 | *Tomentosae* | Diploid | | | 50.4 | 2.7 |
| Nobt | *Nicotiana obtusifolia* | ITB (MPI-ICE) | *Trigonophyllae* | Diploid | | | 17.9 | 1.0 |
| Npal | *Nicotiana palmeri* | Bergerac, 614 | *Trigonophyllae* | Diploid | | | 16.5 | 0.8 |

**Table S1. List and information on the species examined in the study** Information about ploidy and allopolyploids' progenitors are taken from (*34*).

**Table S2**

| | | *Sylvestres* | *Noctiflorae* | *Petunioides* | *undalatae* | *Paniculatae* | *Trigonophyllae* | *Tomentosae* | Age (MYA) |
|---|---|---|---|---|---|---|---|---|---|
| | | **Nsyl** | | **Natt** | | **Npan** | **Nobt** | | |
| *Nicotiana* | **Ntab** | 38.0 | | | | | | Ntom | <0.2 |
| *Rusticae* | **Nrus** | | | | Nund | 36.1 | | | <0.2 |
| *Polydiclae* | **Ncle** | | | 30.6 | | | 32.6 | | ~1 |
| *Repandae* | **Nnes** | 42.9 | | | | | 45.6 | | ~4.5 |
| | **Nnud** | 51.2 | | | | | 43.3 | | ~4.5 |
| | **Nrep** | 41.1 | | | | | 45.4 | | ~4.5 |
| | **Nsto** | 37.2 | | | | | 42.5 | | ~4.5 |
| *Suaevolentes* | **Nben** | 39.3 | ? | ? | | | | | ~10 |
| | **Ngoo** | 40.8 | ? | ? | | | | | ~10 |
| | **Nwut** | 35.5 | ? | ? | | | | | ~10 |

| | | *Sylvestres* | *Noctiflorae* | *Petunioides* | *undalatae* | *Paniculatae* | *Trigonophyllae* | *Tomentosae* | Age (MYA) |
|---|---|---|---|---|---|---|---|---|---|
| | | **Nsyl** | | **Natt** | | **Npan** | **Nobt** | | |
| *Nicotiana* | **Ntab** | 12.0 | | | | | | Ntom | <0.2 |
| *Rusticae* | **Nrus** | | | | Nund | 6.9 | | | <0.2 |
| *Polydiclae* | **Ncle** | | | 8.0 | | | 10.1 | | ~1 |
| *Repandae* | **Nnes** | 10.2 | | | | | 16.8 | | ~4.5 |
| | **Nnud** | 16.4 | | | | | 9.8 | | ~4.5 |
| | **Nrep** | 10.0 | | | | | 17.2 | | ~4.5 |
| | **Nsto** | 10.3 | | | | | 15.2 | | ~4.5 |
| *Suaevolentes* | **Nben** | 10.9 | ? | ? | | | | | ~10 |
| | **Ngoo** | 12.9 | ? | ? | | | | | ~10 |
| | **Nwut** | 11.6 | ? | ? | | | | | ~10 |

**Table S2. Evolutionary distances between allotetraploid species and closest diploid progenitors examined in our study.** Distances were calculated by computing neighbor-joining phylometabolomics tree from Euclidean distances between whole-tissue molecular network (**top table**) and mass motif (**lower table**) representations (See **Figure 2** for the complete tree). Closest progenitors to focal allotretraploids species were not systematically analysed in the study. When not examined, names of these mapped progenitors are provided: Nund, *Nicotiana undulata*; Ntom, *Nicotiana tomentosiformis*. Information about allopolyploid ages and progenitors are taken from (*34*).

**Table S3.**

| Source | Link | Licence |
|---|---|---|
| Natural Products Databse of the UEFS, The State University of Feriera De Santana, Bahia, Brazil( Zinc) | http://zinc.docking.org/ | Whereas you are free to share the results of a ZINC search or a screen of molecules from ZINC, you may not redistribute major portions of ZINC without the express written permission of John Irwin chemistry4biology at gmail.com. |
| Molport Natural Products (Zinc) | http://zinc.docking.org/ | Whereas you are free to share the results of a ZINC search or a screen of molecules from ZINC, you may not redistribute major portions of ZINC without the express written permission of John Irwin chemistry4biology at gmail.com. |
| COCONUT and Natural Products Online | https://coconut.naturalproducts.net/ | If you use data from COCONUT Online, appropriate citation enables readers to locate the original source of the work. |
| AfroDb Natural Products (Zinc) | http://zinc.docking.org/ | Whereas you are free to share the results of a ZINC search or a screen of molecules from ZINC, you may not redistribute major portions of ZINC without the express written permission of John Irwin chemistry4biology at gmail.com. |
| Aster Sunflower family NP (Zinc) | http://zinc.docking.org/ | Whereas you are free to share the results of a ZINC search or a screen of molecules from ZINC, you may not redistribute major portions of ZINC without the express written permission of John Irwin chemistry4biology at gmail.com. |
| Human Metabolome Database Plant (Zinc) | http://zinc.docking.org/ | Whereas you are free to share the results of a ZINC search or a screen of molecules from ZINC, you may not redistribute major portions of ZINC without the express written permission of John Irwin chemistry4biology at gmail.com. |
| Human Metabolome Database Microbe (Zinc) | http://zinc.docking.org/ | Whereas you are free to share the results of a ZINC search or a screen of molecules from ZINC, you may not redistribute major portions of ZINC without the express written permission of John Irwin chemistry4biology at gmail.com. |
| Human Metabolome Database Food (Zinc) | http://zinc.docking.org/ | Whereas you are free to share the results of a ZINC search or a screen of molecules from ZINC, you may not redistribute major portions of ZINC without the express written permission of John Irwin chemistry4biology at gmail.com. |
| Mexican natural products (Zinc) | http://zinc.docking.org/ | Whereas you are free to share the results of a ZINC search or a screen of molecules from ZINC, you may not redistribute major portions of ZINC without the express written permission of John Irwin chemistry4biology at gmail.com. |

| | | |
|---|---|---|
| Knapsack | http://www.knapsackfamily.com | CAUTION: (C) Any content included in KNApSAcK database cannot be re-distributed or used for commercial purposes by any user without contacting with KNApSAcK DB group (skanaya[at]gtc.naist.jp). |
| FooDB (Zinc) | http://zinc.docking.org/ | Whereas you are free to share the results of a ZINC search or a screen of molecules from ZINC, you may not redistribute major portions of ZINC without the express written permission of John Irwin chemistry4biology at gmail.com. |
| The Natural Products Atlas | https://www.npatlas.org/ | Attribution 4.0 International (CC BY 4.0) |
| Biopurify Phytochemicals (Zinc) | http://zinc.docking.org/ | Whereas you are free to share the results of a ZINC search or a screen of molecules from ZINC, you may not redistribute major portions of ZINC without the express written permission of John Irwin chemistry4biology at gmail.com. |
| GOLM | http://gmd.mpimp-golm.mpg.de/ | Academic users may download the material offered on the site for their non-commercial use, but all copyright and other proprietary notices contained in the materials are to be retained. |
| BMRB | https://bmrb.io/ | The data are supplied by the NMR community and are made publicly available free of charge. |
| MetabolomicsWorkbench | https://www.metabolomicsworkbench.org/ | You may download articles and web pages from this site for your personal, non-commercial use only, provided that you keep intact all authorship, copyright and other proprietary notices. |
| UNPD_DB | http://pkuxxj.pku.edu.cn/UNPD | no longer available online |
| Herbal Ingredients In-Vivo Metabolism (Zinc) | http://zinc.docking.org/ | Whereas you are free to share the results of a ZINC search or a screen of molecules from ZINC, you may not redistribute major portions of ZINC without the express written permission of John Irwin chemistry4biology at gmail.com. |
| Nubbe Natural Products (Zinc) | http://zinc.docking.org/ | Whereas you are free to share the results of a ZINC search or a screen of molecules from ZINC, you may not redistribute major portions of ZINC without the express written permission of John Irwin chemistry4biology at gmail.com. |
| NPACT Database (Zinc) | http://zinc.docking.org/ | Whereas you are free to share the results of a ZINC search or a screen of molecules from ZINC, you may not redistribute major portions of ZINC without the express written permission of John Irwin chemistry4biology at gmail.com. |

| | | |
|---|---|---|
| Traditional Chinese Medicine Database@Taiwan (Zinc) | http://zinc.docking.org/ | Whereas you are free to share the results of a ZINC search or a screen of molecules from ZINC, you may not redistribute major portions of ZINC without the express written permission of John Irwin chemistry4biology at gmail.com. |
| Herbal Ingredients Targets (Zinc) | http://zinc.docking.org/ | Whereas you are free to share the results of a ZINC search or a screen of molecules from ZINC, you may not redistribute major portions of ZINC without the express written permission of John Irwin chemistry4biology at gmail.com. |
| Lotus the natural products occurrences database | https://lotus.naturalproducts.net/ | If you use data from LOTUS Online, appropriate citation enables readers to locate the original source of the work. |

**Table S3. List of chemical databases used to construct the 1 million natural product database and predicting *in silico* spectra out of it.** The *in silico* spectra are accessible at Zenodo: https://doi.org/10.5281/zenodo.6536010 . The database was constructed as described in the Supplementary Text "**Creating an *in silico* MS/MS library for approximately 1 million natural products and optimizing its rapid interrogation**".
